# Heavy-tailed connections support robust neural communication

**DOI:** 10.64898/2026.10.21.760834

**Authors:** Jose M. Betancourt, Varun Varanasi, Damon A. Clark, Christopher W. Lynn

## Abstract

In the synaptic wiring between neurons, some rare connections are orders of magnitude stronger than most. This heavy-tailed connectivity is ubiquitous across different species, brain regions, and cell types, suggesting that it serves a general purpose for neural function. Here, we show, both analytically and in simulations, that heavy-tailed connections reduce communication errors in the presence of synaptic noise. In minimal models of individual neurons, noisy connections induce errors at a rate that depends only on a simple network statistic, which we refer to as robustness. For a fixed number of incoming synapses, neurons with broader distributions of connection strength are more robust and, therefore, admit fewer errors. Across several connectomes, we find that real neurons are consistently more robust than comparable null networks. In the whole brain of the fruit fly *Drosophila*, robustness increases with distance from the sensory periphery, such that central neurons are more robust than those involved in sensory processing. Together, these results indicate that heavy-tailed connections may arise universally to fortify the brain against noise in communication.

## Main

Neurons communicate through physical connections composed of synapses.^1^ Competing with many sources of biological noise, signals propagate along synaptic wiring to execute vital neural functions.^2^ The revolution in connectomics now provides maps of these connections across multiple species, from *C. elegans*,^3, 4^ mice,^5^ and zebrafish^6^ to the entire brain of the fruit fly *Drosophila*.^7–10^ Universally, these networks share a defining feature: connection strengths are heavy-tailed.^11, 12^ While most neurons interact only weakly, a handful of important connections are orders of magnitude stronger, forming the backbone of neural circuitry. Yet despite this ubiquity, the advantage of heavy-tailed connections for neural function remains unclear.

Take the complete adult brain of the female fruit fly, which consists of nearly 140,000 neurons, each with an average of 109 incoming connections (Table 1).^7, 8^ In invertebrates like the fly, connections typically grow stronger by increasing the number of synapses, such that synapse count approximates functional connection strength.^13−15^ While the average connection comprises only 3-4 synapses, some specific connections consist of over 2,000 individual synapses (Fig. 1a). This heavy tail in the distribution of connection strength is highly non-random. Shuffling the placement of synapses destroys the strong connections, even after accounting for the variability between neurons (Fig. 1a). Heavy-tailed connectivity is not limited to a single neuron type or brain region, but instead arises consistently across the entire brain (Fig. 1b). Similar distributions appear in the combined brain and nerve cord of the female fly (Fig. 1c),^9^ the nerve cord of the male fly (Fig. 1d),^10^ the nematode *C. elegans* (Fig. 1e),^3, 4^ and the mouse retina (Fig. 1f),^5^ among other connectomes.^6, 11^ Even in measurements of activity (rather than structure), some rare pairs of neurons are correlated much more strongly than most.^12, 16−19^ Given the universality of heavy-tailed connections, does there exist an equally general purpose for neural function?

**Table 1.**
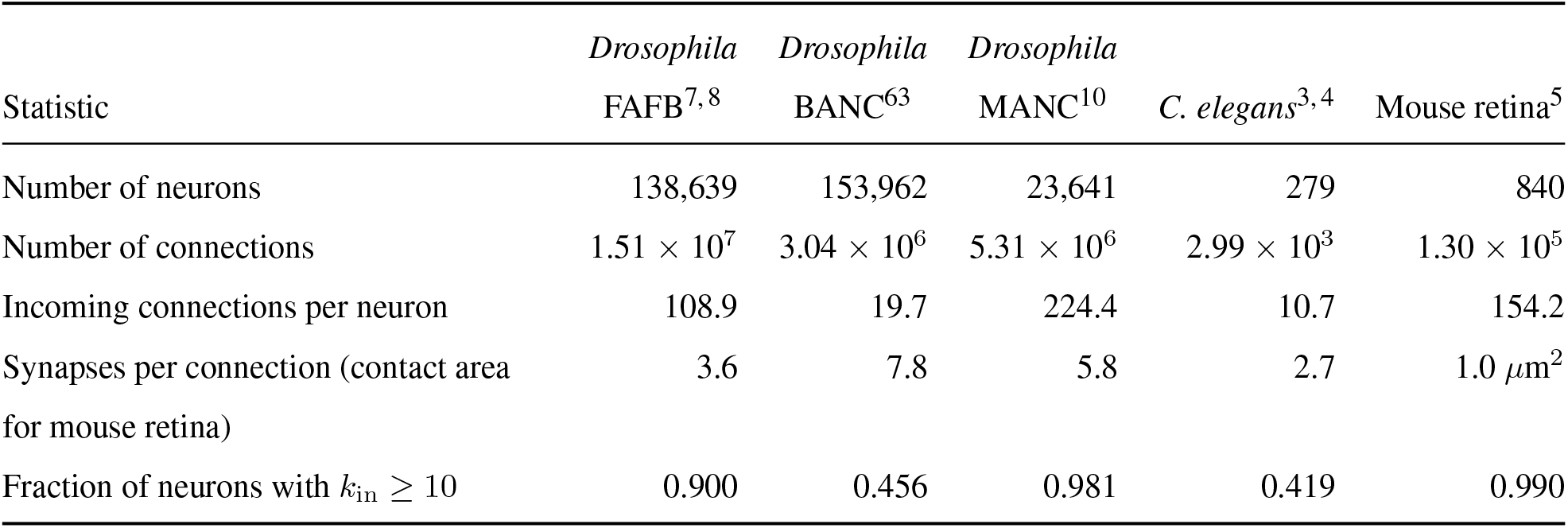
Statistics of neuronal connectomes.

**Fig. 1.**
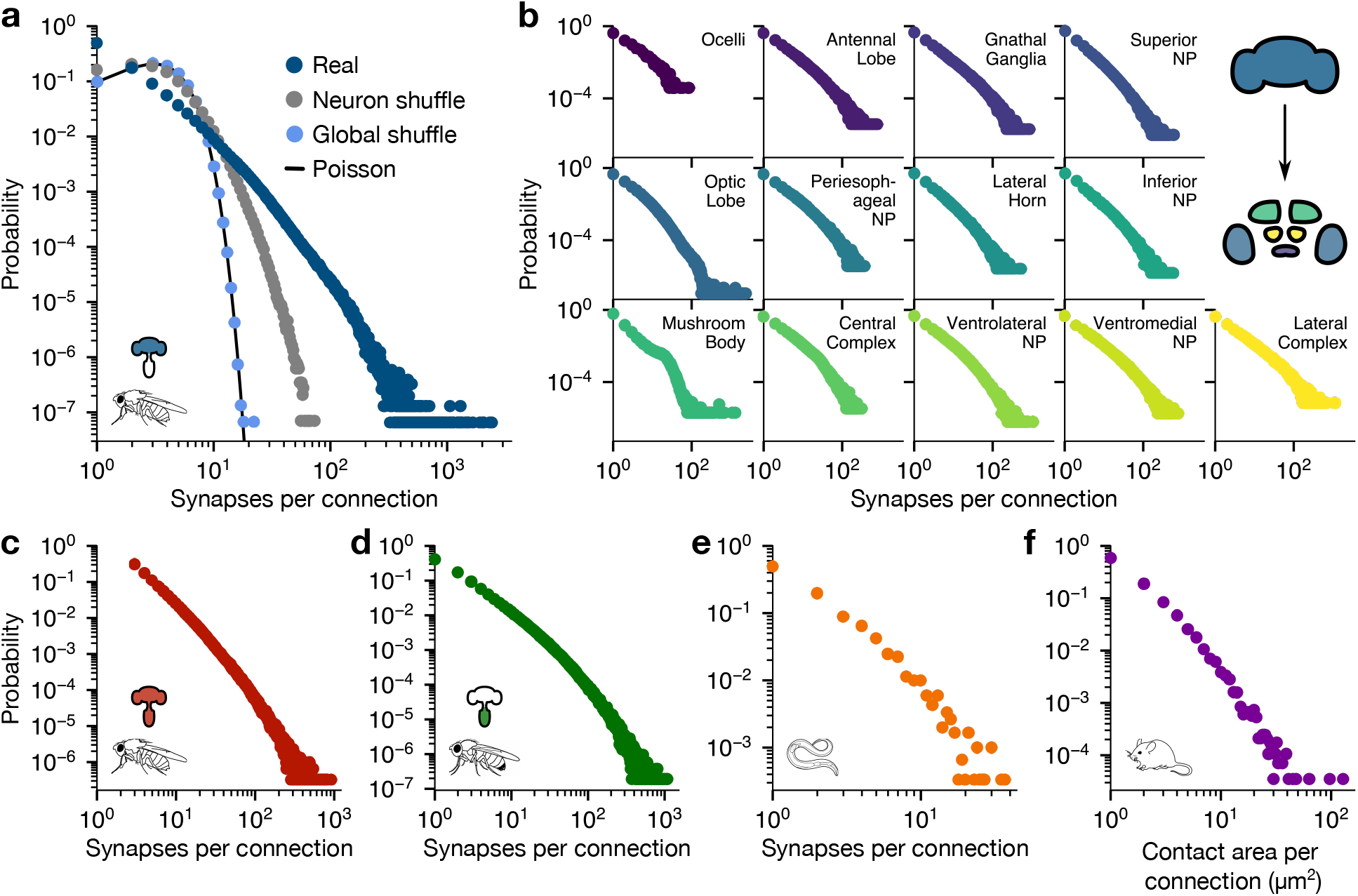
Heavy-tailed synaptic connectivity across species and systems. **a**, Distribution of synapses per connection between neurons in the whole brain of the female fruit fly *Drosophila*^7,8^ (dark blue), after shuffling the synapses among incoming connections to each neuron (grey) and after shuffling synapses among all connections in the network (light blue); solid line indicates a Poisson distribution. **b**, Distributions of synapses per connection for different regions of the female fruit fly brain.^7,8^ Connections are assigned to regions based on the plurality of synapses (Methods). **c**-**e**, Distributions of the number of synapses per connection in the combined brain and nerve cord of a *Drosophila* female (**c**),^9^ the *Drosophila* male nerve cord (**d**),^10^ and the nematode *C. elegans* (**e**).^3,4^ **f**, Distribution of the total contact area (rounded to the nearest *µ*m^2^) per connection between neurons in the inner plexiform layer of the mouse retina.^5^

### Noisy connections induce errors in communication

Unlike artificial networks, in which weights are fixed with arbitrary precision, communication between neurons must contend with random fluctuations across many scales. Two neurons performing the same function in genetically identical organisms, or within the same organism, form different connections.^20−22^ Over time, individual neurons create new synapses, prune old synapses, and strengthen connections via plasticity.^23−25^ Even with connectivity held fixed, the response of a neuron can change due to stochastic ion channels,^26^ variable neurotransmitter release,^27, 28^ and thermal noise.^2, 29^ Thus, in the language of information theory, synaptic connections function as noisy communication channels between upstream inputs and a downstream output.^30^ Yet in the face of significant synaptic noise, key neural functions remain intact.^2, 31−33^

To investigate how synaptic noise impacts communication, consider a minimal McCulloch-Pitts (MP) neuron or perceptron (Fig. 2a),^34, 35^ which receives inputs from *n* presynaptic neurons (*i* = 1, …, *n*) that are either active (*x*_*i*_ = 1) or silent (*x*_*i*_ = −1). These inputs are weighted by connection strengths *w*_*i*_, combined with a bias *b*, and then passed through a threshold to produce the output *y* = sgn ( ∑*i w*_*i*_*x*_*i*_ + *b*). Although a simplification of real neurons, this model captures key features of neural interactions and forms the basis for artificial networks.^19, 36−38^ Biological noise leads to random fluctuations *ŵ*_*i*_ in the weights, such that the instantaneous strength of a connection is given by 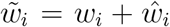 (Fig. 2a). For example, if each unit of connection strength (e.g., each synapse or unit of contact area) is subject to independent noise of strength *σ*, then connections fluctuate according to Var(*ŵ*_*i*_) = *σ*^2^*w*_*i*_. In turn, these fluctuations produce errors in communication: for the same inputs ***x*** = {*x*_*i*_}, the postsynaptic neuron may generate a different output 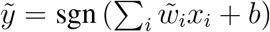. Geometrically, the original neuron with weights ***w*** = {*w*_*i*_} defines a decision boundary in the space of inputs (Fig. 2b).^39^ Noise in the weights shifts this boundary, and any inputs ***x*** that cross the hyperplane result in errors 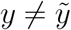. Across all inputs and fluctuations, the neuron admits errors at a rate 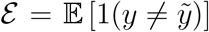, where 1(·) is the indicator function and E[·] represents an average over both inputs and noise. Importantly, the error rate depends only on the nature of the communication channel itself, which is determined by the strengths of connections.

**Fig. 2.**
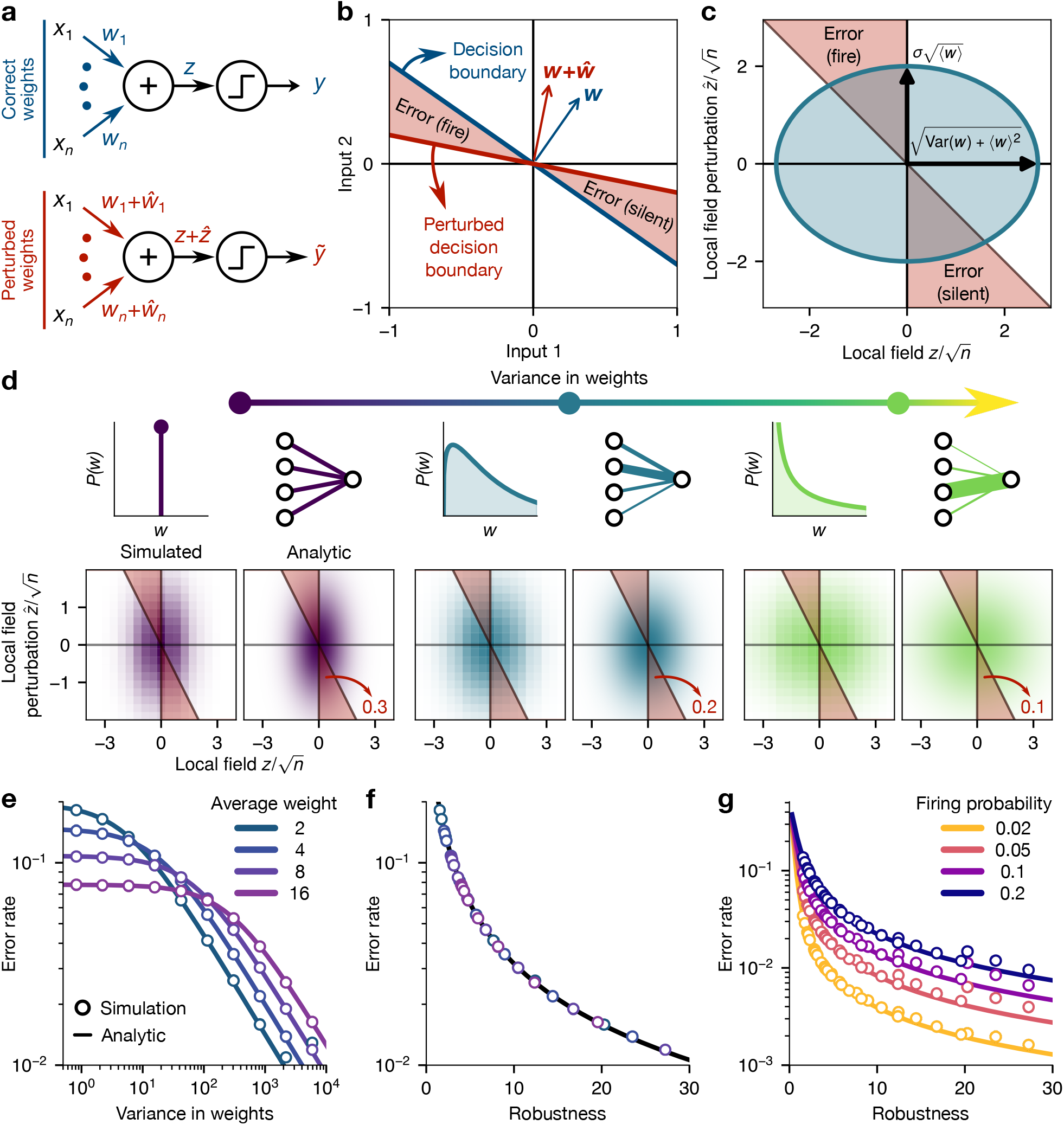
Quantifying errors due to synaptic noise. **a**, Minimal model of noise in neural communication. *Top:* In a McCulloch-Pitts (MP) neuron or perceptron, the neuron passes *n* inputs *x*_*i*_ through linear weights *w*_*i*_ to form a local field *z*, which determines the binary output *y* = sgn(*z*). *Bottom:* Noise *ŵ*_*i*_ in the weights yields a perturbed local field 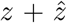, which can lead to a different output 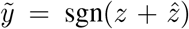. **b**, Synthetic neurons define decision planes in the high-dimensional input space. Noise in the weights shifts the boundary, and inputs that cross the hyperplane represent errors 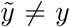. **c**, Joint distribution of the local field *z* and perturbation 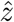 determines the error rate. For independent inputs, as *n* → ∞ the distribution approaches a Gaussian with widths defined by the mean and variance of the weights (Methods). Blue region illustrates a level set of the distribution with *b* = 0 and E(*x*_*i*_) = 0, and arrows represent standard deviations. **d**, Simulated and analytic distributions of local fields *z* and perturbations 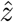 for three neurons with *n* = 10^3^ inputs and noise of strength *σ* = 1. From left to right, incoming weights are drawn from distributions with equal mean and increasing variance, leading to decreasing errors (Methods). Color is proportional to the probability density. Inset numbers indicate error rates. **e**, Error rates for synthetic neurons with fixed average connection strength and increasing variance. **f**, Error rate as a function of robustness *r*(***w***) for the same neurons as panel **e. g**, Error rate as a function of robustness for neurons with the same weights as panels **e** and **f**, but different firing probabilities (Methods). In **e-g**, each neuron has *n* = 10^3^ inputs, noise of strength *σ* = 1, and weights drawn from gamma distributions with fixed mean and variance. Lines are analytic predictions and points are averages over simulations of 10^3^ neurons (Methods).

To understand how connections shape errors, an important quantity is the local field *z* = ∑*i w*_*i*_*x*_*i*_ + *b*, which dictates whether the neuron fires (*z >* 0) or not (*z <* 0).^40^ To produce an error, the perturbation to the local field 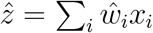 must be large enough to change the sign of *z* (Fig. 2c). In the simplest scenario, the bias *b* is zero and the inputs are independent and symmetric (E(*x*_*i*_) = 0). As the number of inputs *n* grows, the distributions of local fields *z* and perturbations 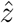 are defined exactly by the mean ⟨*w*⟩ and variance Var(*w*) of the weights (Fig. 2c; Methods). Increasing the variance pushes the local field away from the decision boundary (*z* = 0), such that errors require stronger perturbations. Thus, for fixed average strength ⟨*w*⟩, as connections become more heterogeneous, the neuron admits fewer errors (Fig. 2d). In the limit *n* → ∞, we arrive at an analytic prediction for the error rate,

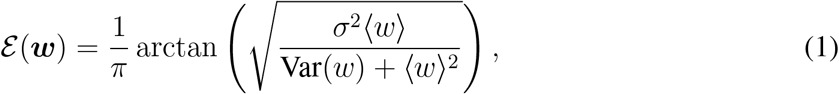

which we confirm in simulations (Fig. 2e). The error rate depends non-monotonically on the average strength ⟨*w*⟩: for low variance Var(*w*), errors are minimized by strong connections, while for high variance, weak connections minimize errors. By contrast, for any average strength ⟨*w*⟩, increasing the variance leads to a monotonic decrease in errors (Fig. 2e).

Strikingly, Eq. (1) reveals that errors do not depend independently on the mean and variance in the weights, but instead through a single statistic 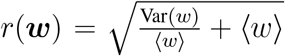. Since 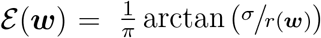, the statistic *r*(***w***) defines the strength of noise *σ* that a neuron can withstand before errors become significant; that is, it defines the neuron’s *robustness*. When *r*(***w***) ≫ *σ*, errors vanish (*ε* → 0), and when *r*(***w***) ≪ *σ*, errors dominate (*ε* → 1*/*2). Plotting the error rate as a function of robustness, we find that all of the data in Fig. 2e collapse onto a single analytic curve (Fig. 2f). Even without many of the simplifying assumptions, the same structure remains. As long as the average output E(*y*) is held fixed, the error rate still decreases monotonically with robustness for arbitrary bias *b* and input averages E(*x*_*i*_) (Fig. 2g; Methods). One can also include correlations between inputs, continuous inputs and outputs, and different noise structures Var(*ŵ*_*i*_) (Supplementary Information). In all cases, increasing the variance in connection strengths pushes the neuron away from the decision boundary, thereby reducing errors in communication.

### Heavy-tailed connectivity supports robust communication

Ultimately, we seek to understand how real distributions of connection strength impact communication (Fig. 1). Across the fruit fly brain, neurons are highly heterogeneous, varying by over three orders of magnitude in the number of inputs *n* (Fig. 3a), one order of magnitude in average connection strength ⟨*w*⟩ (Fig. 3b), and four orders of magnitude in variance Var(*w*) (Fig. 3c).^7, 8^ For each neuron, we construct a synthetic MP neuron with weights *w*_*i*_ defined by the number of synapses in each incoming connection (Fig. 2a), which is a reasonable approximation in the case of the fruit fly.^13−15^ We then simulate activity before and after adding noise to every synapse. Due to the heterogeneity between neurons, the same amount of synaptic noise induces a wide range of errors; yet despite this variability, errors are consistently predicted by the analytic rate in Eq. (1) (Fig. 3d). Grouping neurons based on their average connection strength ⟨*w*⟩, we find that errors systematically decrease with the variance in weights (Fig. 3e). Moreover, these errors are completely characterized by the robustness of each neuron *r*(***w***), such that over 10^5^ neurons across the entire brain collapse to one curve (Fig. 3f). This collapse—and our analytic predictions—hold consistently for different firing rates (Fig. 3g), amounts of synaptic noise (Fig. 3h), and definitions of the weights *w*_*i*_ (Supplementary Information).

**Fig. 3.**
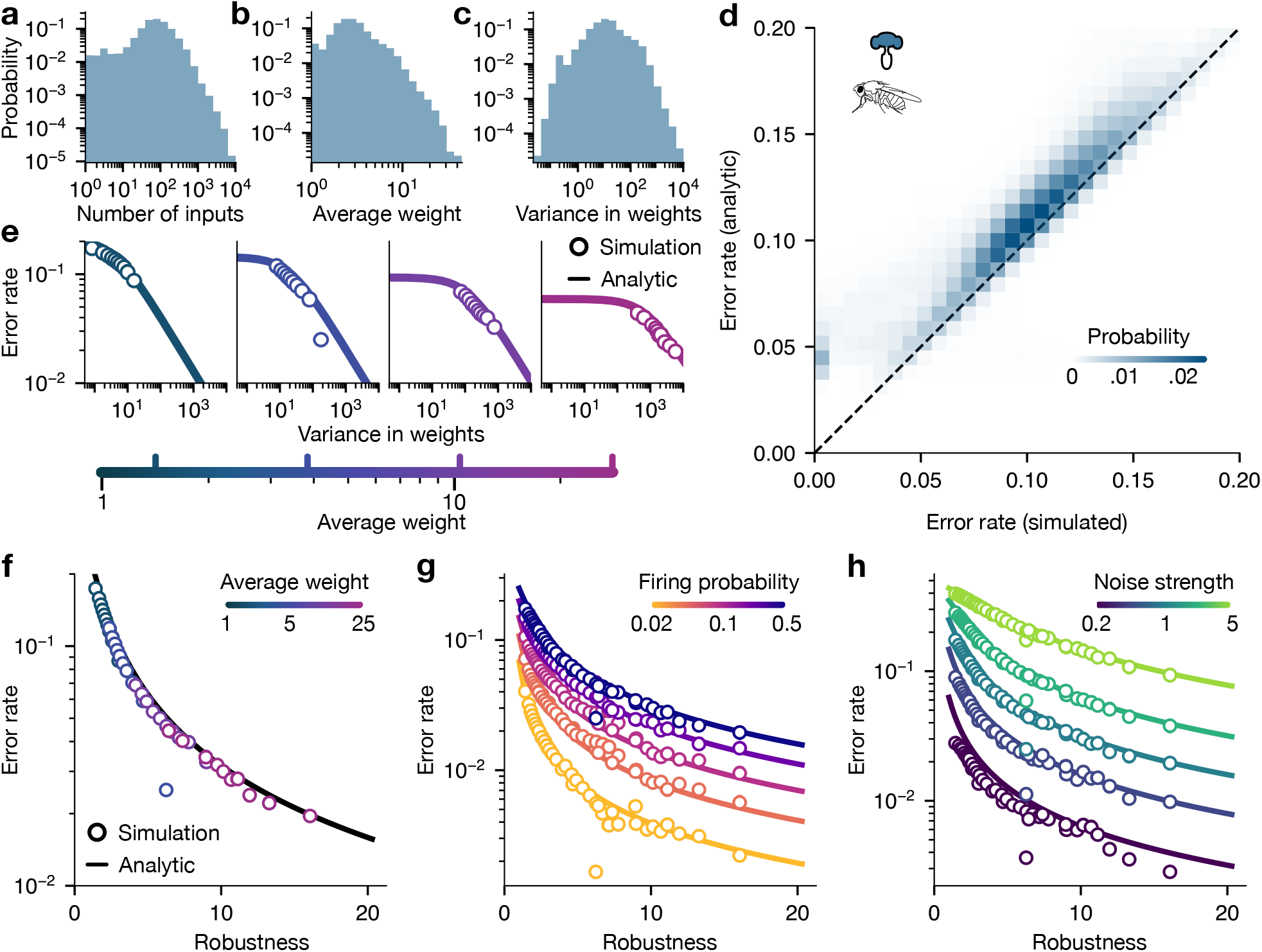
Communication errors with real connection strengths. **a-c**, Distributions of numbers of inputs *n* (**a**), average connection strength ⟨*w*⟩ (**b**), and variance in connection strengths Var(*w*) (**c**) across neurons in the whole brain of the female fruit fly.^7,8^ **d**, For each neuron in the fly brain, we construct a synthetic neuron with weights defined by the number of synapses with presynaptic partners. Distribution of simulated and analytic error rates across all neurons with at least ten inputs. Unless otherwise specified, bias is zero (*b* = 0), inputs are independent and symmetric (E(*x*_*i*_) = 0), and synaptic noise is applied with strength *σ* = 1. **e**, Error rates as a function of variance in connection strengths for increasing average strength. Neurons are grouped by the mean and variance of their incoming connection strengths (Methods). Points represent the median values within each group, and solid lines are analytic predictions from Eq. (1). **f**, Error rate as a function of robustness *r*(***w***) for the groups of neurons in **e. g-h**, Error rate as a function of robustness for the same groups with different firing rates (**g**) and different levels of synaptic noise (**h**). See Methods for simulation details.

Across different organisms and brain regions, the fact that connections are heavy-tailed suggests that neuronal communication may be highly robust to synaptic noise (Fig. 1).^11^ To test this hypothesis, a natural comparison is a null neuron with the same total number of inputs and synapses (such that the average connection strength ⟨*w*⟩ is fixed), but with synapses shuffled among connections. These null neurons have Poisson distributions of connection strength, characterized by Var(*w*) = ⟨*w*⟩. In the fly brain, we see that most neurons have variance larger than Poisson, some by multiple orders of magnitude (Fig. 4a, *inset*).^7, 8^ These highly variable weights, punctuated by abnormally strong connections, make the vast majority of neurons (93%) more robust than their shuffled counterparts (Fig. 4a).

**Fig. 4.**
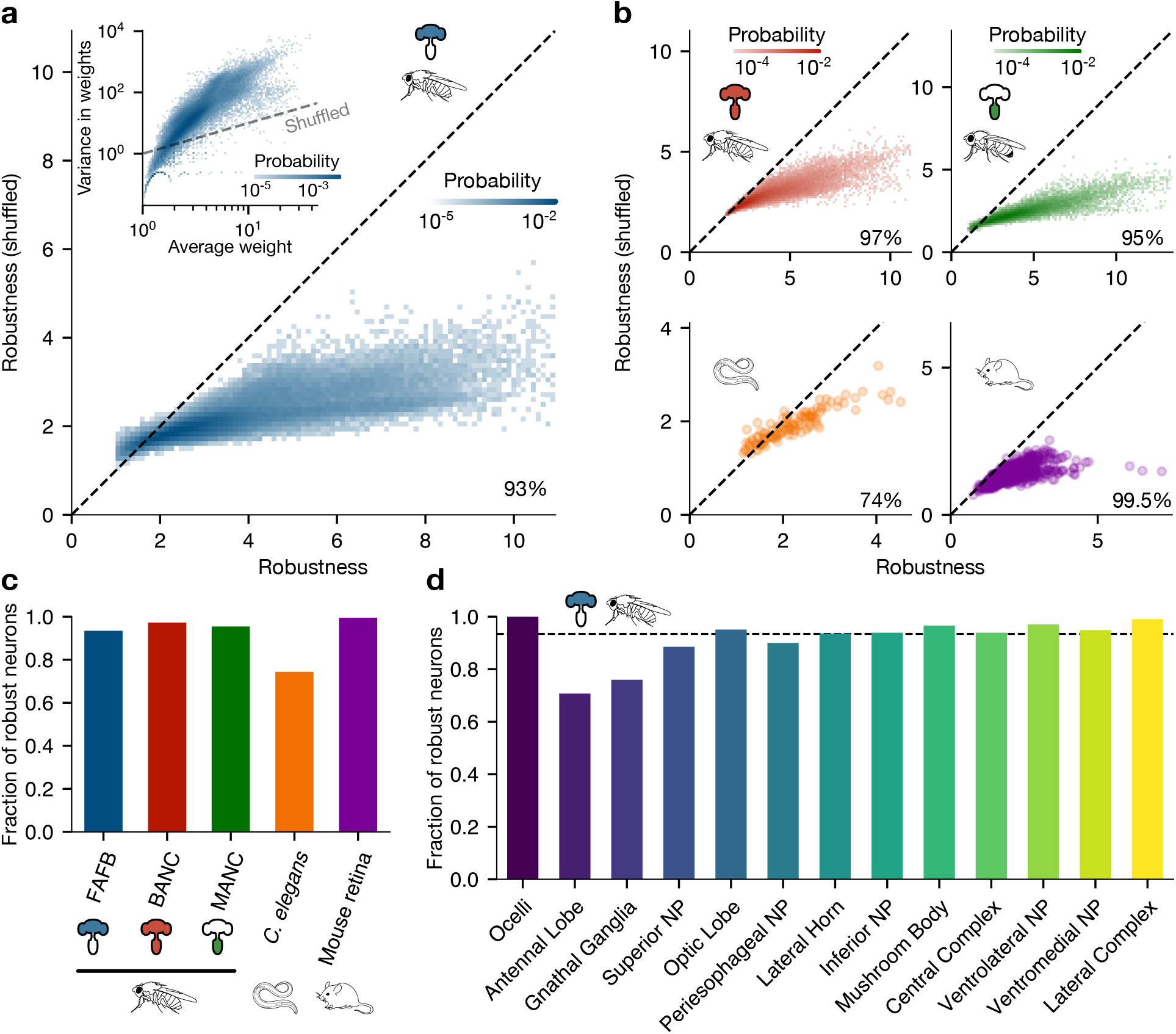
Robust communication driven by heavy-tailed connectivity. **a**, Distribution of robustness *r*(***w***) across neurons in the female fruit fly brain^7,8^ before and after shuffling synapses among incoming connections. Percentage indicates fraction of neurons below the dashed line, which are more robust than shuffled nulls. *Inset:* Distribution of average connection strength ⟨*w*⟩ and variance in strengths Var(*w*) over neurons in the female fly brain,^7,8^ where each weight represents the number of synapses in an incoming connection. Dashed line represents the Poisson variance for shuffled neurons. **b**, Distribution of robustness before and after shuffling connection strength among incoming connections for neurons in the combined brain and nerve cord of a *Drosophila* female (*top left*),^9^ the nerve cord of a *Drosophila* male (*top right*),^10^ the nematode *C. elegans* (*bottom left*),^3,4^ and the inner plexiform layer of the mouse retina (*bottom right*).^5^ Dashed lines represent equality, and percentages indicate the fraction of neurons with higher robustness than shuffled nulls (Methods). **c**, Fractions of neurons with robustness larger than shuffled counterparts in different connectomes. **d**, Fractions of neurons with robustness larger than shuffled nulls in different regions of the female fruit fly brain.^7,8^ Dashed line represents the percentage for the whole brain. Neurons are assigned to brain regions based on the plurality of incoming synapses (Methods). In all connectomes, we only study neurons with at least ten incoming connections.

We repeat the same analysis for many neurons spanning different individuals, systems, and species. In the combined brain and nerve cord of the female fruit fly, a staggering 97% of neurons are more robust to noise than comparable nulls (Fig. 4b).^9^ This percentage drops slightly in the nerve cord of the male fly (95%)^10^ and more dramatically in *C. elegans* (74%; Fig. 4b),^3, 4^ reflecting limitations on very strong connections in the nematode (Fig. 1e).^11^ In vertebrates, connections are strengthened by increasing not only the number of synapses, but also the sizes of individual synapses.^41, 42^ Thus, for neurons in the mouse retina, we construct synthetic neurons with weights *w*_*i*_ defined by the total contact area with each partner (Fig. 1f).^5^ After shuffling contact area among connections, thereby destroying the heavy-tailed connectivity, we find that 99.5% of neurons become less robust to noise (Fig. 4b). Even within a single organism, across all regions of the fly brain, the majority of neurons are highly robust (Fig. 4d).^7, 8, 43^ Together, these results suggest that heavy-tailed connectivity, which is observed universally in biological networks, supports reliable communication in the face of synaptic noise.

### Robustness across an adult brain

In the fruit fly, we have seen that robustness varies dramatically from one neuron to another (Fig. 4a-b).^7−10^ Is this robustness distributed evenly throughout the brain, or instead, can some regions or cell types withstand more noise than others? To compare across cells, we need a standardized scale that accounts for differences in average connection strength ⟨*w*⟩. For any neuron, robustness is minimized by spreading synapses uniformly among inputs, yielding 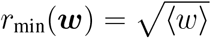 By distributing synapses randomly, robustness increases to 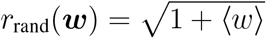. This difference defines a natural scale, leading to a normalized robustness,

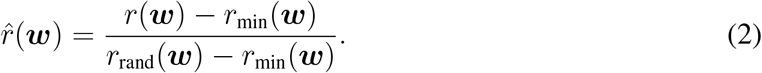

Any normalized robustness above (below) one means that synapses are distributed to withstand more (less) noise than random. Indeed, after shuffling synapses among connections, we see that normalized robustness peaks sharply at 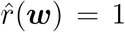 (Fig. 5a). By contrast, real neurons exhibit a much wider range of robustness, with the median neuron 3.4 times more robust than random.

**Fig. 5.**
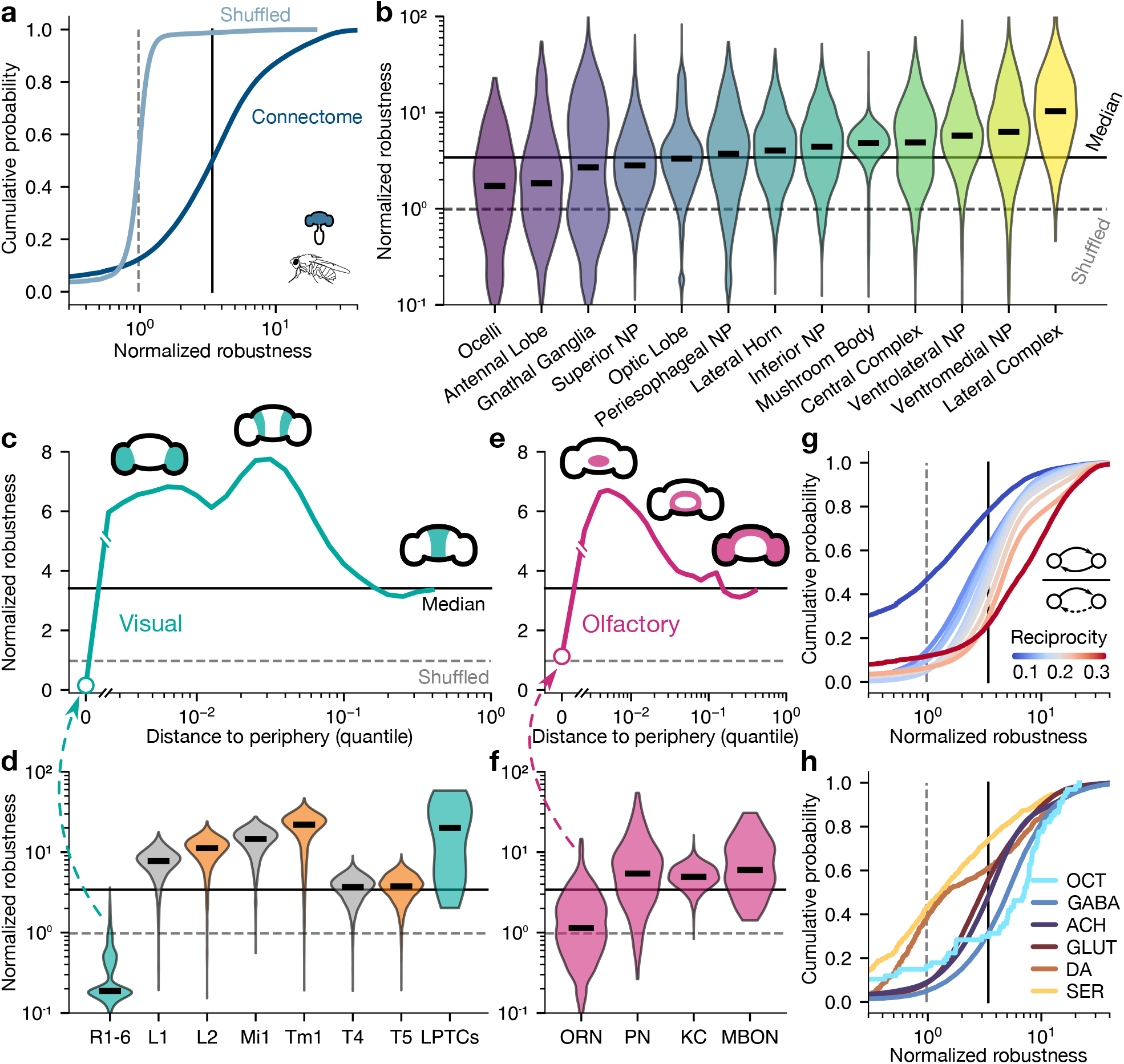
Robustness in the *Drosophila* brain. **a**, Cumulative distributions of normalized robustness across neurons in the fruit fly brain before and after shuffling synapses among incoming connections.^7,8^ Lines indicate medians. **b**, Distribution of normalized robustness across neurons within each brain region. Neurons are assigned to brain regions based on the plurality of their incoming synapses (Methods). **c**, Normalized robustness versus distance from the visual periphery, calculated using a spreading process initialized at the photoreceptors R1-6 (Methods).^7,8,^44 Values represent medians among neurons grouped by their (log) quantile distance to the periphery. **d**, Distribution of normalized robustness for specific cell types along the ON (teal and grey) and OFF (teal and orange) motion pathways in the visual system.^45,46^ LPTCs include HS, VS, H1/H2, and DCH/VCH neurons. **e**, Normalized robustness versus quantile distance from the olfactory periphery, calculated using a spreading process initialized at the olfactory receptor neurons (ORNs). **f**, Distribution of normalized robustness for specific cell types along the odor encoding pathway.^47^ **g**, Distributions of normalized robustness for neurons grouped by reciprocity, or the fraction of neighbors that are both pre- and postsynaptic partners (Methods).^48^ **h**, Distributions of normalized robustness for neurons grouped by neurotransmitter release (Methods). In all panels, dark solid lines represent the median normalized robustness across the entire fruit fly brain, and light dashed lines represent neurons with shuffled synapses.

The fly brain naturally partitions into distinct anatomical regions, which are responsible for different neural functions.^7, 8, 43^ Because neurons can have incoming or outgoing synapses in different brain regions, we assign each neuron to the region that contains the most of its incoming synapses (Methods). Between regions, we find that robustness varies significantly, increasing by an order of magnitude from the least robust region (ocelli) to the most robust (lateral complex; Fig. 5b). Notably, a hierarchy emerges in which regions involved in sensory processing (e.g., antennal and optic lobes)^49, 50^ are less robust than those associated with central functions (e.g., mushroom body and central complex).^51, 52^

To understand whether robustness increases from the sensory periphery to the central brain, we define the distance of each neuron from an initial set of sensory cells using a spreading process (Methods).^7, 8, 44^ For example, vision begins at the photoreceptors R1-8,^53^ which, due to their small number of weak synaptic inputs, have normalized robustness near zero (Fig. 5c,d). As we move away from the periphery, robustness increases dramatically to over seven times higher than random, before returning to the whole-brain median (Fig. 5c). We observe a similar pattern in specific visual circuits, such as the ON motion pathway, which detects increases in light intensity.^45, 46^ Robustness rises abruptly from the photoreceptors R1-6 to downstream interneurons L1 and Mi1, and decreases to the median at the outputs T4 (Fig. 5d; *grey*). The OFF pathway, which detects decreases in light intensity, displays a nearly identical pattern (Fig. 5d; *orange*). Comparatively, olfaction begins at receptor neurons (ORNs), which have normalized robustness near one, equivalent to shuffled synapses (Fig. 5e,f). As signals propagate from the periphery, neurons again become multiple times more robust before settling to the median (Fig. 5e). Along the odor encoding pathway,^47^ robustness increases sharply from ORNs to projection neurons (PNs), Kenyon cells (KCs), and the eventual outputs in the mushroom body (MBONs; Fig. 5f). Thus, in both vision and olfaction, neurons become more robust with distance from the sensory periphery.

From the periphery to the central brain, circuits transition from feedforward to highly recurrent, suggesting that robustness may be linked to network structure.^48, 54^ Locally, recurrence is quantified by a neuron’s reciprocity: the fraction of neighbors that are both pre- and postsynaptic partners.^48^ Across the fly brain, as neurons become less feedforward and more reciprocal, they consistently become more robust (Fig. 5g), reinforcing the association between central processing and robustness to synaptic noise. Finally, in addition to anatomical structure and network topology, neuronal interactions are driven by the identity of the neurotransmitter released by each neuron.^7, 8, 55^ We find that neurons involved in both fast excitation and inhibition—releasing GABA, acetylcholine (ACH), and glutamate (GLUT)—all exhibit similar distributions of robustness (Fig. 5h).^55^ By contrast, modulatory neurons that release serotonin (SER) and dopamine (DA) are significantly less robust, while those that release octopamine (OCT)—which mediates memory, arousal, and behavioral state—are more robust.^56−58^ Thus, the significant variation in robustness between neurons is not random, but instead couples tightly to neural function and structure.

## Discussion

Connections between neurons are universally heavy-tailed (Fig. 1),^3−11^ hinting at a fundamental role for neural function. Here we show that in the presence of noise, heavy-tailed connections reduce errors in communication. In synthetic neurons, a single network statistic (robustness) sets the scale at which noise leads to errors (Fig. 2). Even for the same average connection strength, as distributions grow wider, neurons become more robust to noise, thereby suffering fewer errors. Leveraging advances in high-throughput connectomics, we investigate the robustness of hundreds of thousands of neurons spanning several species and neural systems (Figs. 3-5). By concentrating synapses among strong connections, most neurons are highly robust, suggesting that connectivity may be optimized to overcome biological noise.

Our results and their limitations point to concrete future directions. To make analytic progress [Eq. (1)], we employ synthetic models of MP neurons or perceptrons (Fig. 2a). While these models capture key features of neural interactions and have been used to predict neural activity,^19, 36−38, 59^ real neurons involve complex biophysical processes that induce interactions between inputs.^60, 61^ Building upon our framework, one can study robustness in more realistic models of neural activity. This will shed light on how robust communication is shaped not only by synapse count and strength, but also by synapse location, neuron morphology, and other biophysical parameters. Importantly, as long as noise has a sub-multiplicative impact on a neuron’s local field, we still expect heavy-tailed connectivity to reduce errors in communication (Supplementary Information).

Although we focus on individual neurons as stochastic communication channels, signals in the brain propagate through complex networks that involve many neurons.^54^ By wiring neurons together and simulating collective dynamics, one can investigate how synaptic noise leads to errors at the network level.^2^ In turn, this will reveal how different circuit structures support robust communication and optimize information flow.^2, 54^ Finally, our results indicate that distinct parts of the brain are structured to withstand different amounts of noise (Fig. 5), yet the reasons remain unclear. Sensory neurons may be optimized for sensitivity to small changes in inputs at the expense of robustness.^62^ At the same time, signaling in the central brain may be forced to contend with greater noise than at the periphery.^2^ As experiments continue to advance, identifying universal features of network structure and understanding how they support neural functions remain critical outstanding challenges.

## Methods

### Connectome data

The datasets used in this paper were previously published and are openly accessible.^3–5,7–10^ All connectomes were sectioned and imaged using electron microscopy. Identification and classification of neurons and synapses were achieved via a combination of manual labeling and automated processing with subsequent proofreading. We provide the preprocessing code required to reproduce the results from the original datasets (Data Availability). Connections in the mouse retina correspond to contact area between neurons, and are undirected.^5^ For our analysis, we treat each contact as an incoming connection to both neurons involved. The connectome for the female adult fly brain (FAFB) identifies each synapse with pre- and postsynaptic neurons as well as the neuropil in which it is located, which are grouped into brain regions. ^7, 8, 43^ We identify connections with brain regions based on which region contains the plurality of their synapses (for Fig. 1b) and analogously assign neurons based on plurality of their post-synapses (for Figs. 4d and 5b). Roughly 95% of connections consist of synapses within a single brain region, and 70% of neurons have all incoming synapses in a single brain region (Supplementary Information). We assign neurotransmitters to neurons following the FAFB classification.^7,8^

### Simulated error rate

Unless otherwise stated, simulations use zero bias (*b* = 0), inputs *x*_*i*_ are drawn independently and uniformly from {−1, +1}, weight perturbations *ŵ*_*i*_ are drawn from a zero-mean Gaussian with noise strength parameter *σ* = 1, and the error rate for each neuron is averaged over all combinations of 10^3^ input samples and 10^3^ weight perturbations. All synthetic neurons have *n* = 10^3^ inputs. In Fig. 2d, we generate three neurons: one with uniform weights (*w*_*i*_ = 1) and two with gamma-distributed weights of mean ⟨*w*⟩ = 1 and variances Var(*w*) = 0.8 and 3. The histograms use the data from all combinations of 5 × 10^3^ input samples and 5 × 10^3^ weight perturbations. Each data point in Figs. 2e-g represents an average over 10^3^ neurons with gamma-distributed weights with a specified mean and variance. In Fig. 3h, we adjust the noise strength *σ*. In Figs. 2g and 3g, we change the firing rate by tuning the bias *b* using the Gaussian approximation in Eq. (3). In the Supplementary Information, we investigate the effects of sparse inputs [E(*x*_*i*_) *<* 0], correlations between inputs, different noise structures, and different definitions of connection strength.

### Analytic error rate

To derive the analytic prediction for the error rate, we study the limit of many inputs *n* → ∞. For a set of weights ***w*** = {*w*_*i*_} and bias *b*, the local field is *z* = ∑_*i*_ *w*_*i*_*x*_*i*_ + *b*. Given noise ***ŵ*** = {*ŵ*_*i*_}, the perturbed weights 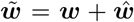 yield a perturbed local field 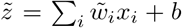. Errors occur when the signs of *z* and 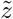 differ, such that the error rate is given by 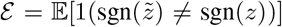, where E(·) is an expectation over realizations of the inputs ***x*** and noise ***ŵ***.

To study the *n* → ∞ limit, we assume that weights *w*_*i*_ are drawn independently from a distribution *P* (*w*) with finite mean and variance; here, ⟨·⟩ denotes an average over *P* (*w*). We assume the inputs *x*_*i*_ are independent and identically distributed for each *n*; this distribution can change as *n* grows, but must converge. We also allow the bias *b* to change. As *n* increases, we hold the output firing probability fixed to a finite value *p*_*f*_ . For large *n*, the central limit theorem dictates that

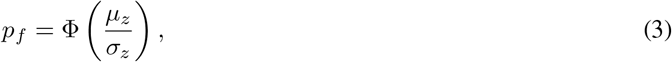

where Φ(·) is the standard normal cumulative distribution function and *µ*_*z*_ and 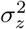 are the mean and variance of *z*, respectively. Moreover, the correlation between *z* and 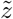 is given by

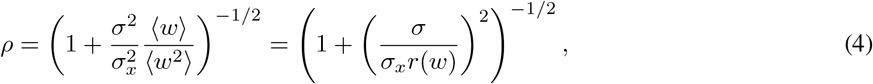

where 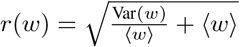 is the robustness of the neuron and 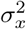 is the input variance.

The distribution of normalized local fields 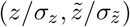 converges to a Gaussian with means 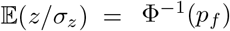 and 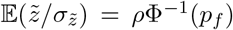, unit variance, and covariance *ρ*. Computing the error rate amounts to computing the mass in the upper-left and lower-right orthants. The solution to the Gaussian orthant problem yields

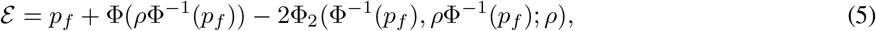

where Φ_2_ is the bivariate standard normal cumulative distribution function (Supplementary Information). We have therefore established that the error rate depends only on the output firing probability *p*_*f*_ and the correlation between local fields *ρ*, which itself depends on the robustness *r*(*w*), noise *σ*, and input variability *σ*_*x*_. Thus, the weights *w*_*i*_ only enter the error rate through the firing probability and robustness. For a firing rate *p*_*f*_ = 1*/*2, the error rate simplifies significantly to Eq. (1).^64^

### Grouping neurons by connectivity statistics

In Figs. 3e−h, we group neurons based on the statistics of their incoming connections. For each neuron, we compute the average number of synapses and the variance in the number of synapses among incoming connections. Since both average strength and variance in strength are approximately log-normally distributed (Figs. 3b-c), we first group neurons into uniformly log-spaced bins by average strength. Then within each bin, we group neurons into deciles by variance in strength. Each point in Figs. 3e-h represents the median over neurons in one group.

### Shuffled neurons

We generate one shuffled neuron for each real neuron with at least ten incoming connections (Fig. 4). For the female fruit fly brain,^7, 8^ fruit fly male nerve cord,^10^ and nematode *C. elegans*,^3, 4^ we randomly shuffle synapses among incoming connections. The fruit fly brain and nerve cord connectome only includes connections with at least three synapses.^9^ To generate shuffled neurons, we first assign three synapses to each incoming connection and then place the remaining synapses randomly. For the mouse retina,^5^ connection strength is functionalized as the total contact area across all contacts comprising one connection, such that the measure of connection strength is no longer discrete. To generate shuffled neurons, we sample uniformly from the set of connection strength vectors ***w*** with total contact area ∑_*i*_ *w*_*i*_ matching the original neuron. See the Supplementary Information for alternative shuffling choices.

### Distance to periphery

For each neuron in the female fruit fly brain, to define the distance from a set of sensory neurons (Fig. 5c,e), we use a network spreading process introduced in Ref. 44. The process begins by defining a seed set of neurons. At each time step, downstream neurons are probabilistically added to the set. Specifically, each neuron *i* in the set succeeds in adding each outside neuron *j* with probability 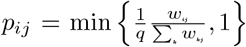, where *w*_*ij*_ is the number of synapses from *i* to *j* and *q* = 0.3 is a threshold.^44^ For each seed set, we simulate the spreading process 10^3^ times. Simulations are terminated when all neurons in the brain are reached, or if no neurons are added in ten consecutive steps. Starting from the visual periphery, the process reaches an average of 98.9% of neurons, while an average of 98.6% are reached when starting from the olfactory periphery. We define the distance from neuron *i* to the periphery as the average number of simulation steps required to add *i* to the set. We then group neurons into quantiles of distance to the periphery.

### Reciprocity

For a given neuron *i*, we define 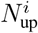 and 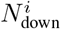 to be the set of neurons immediately upstream or down-stream of *i*, respectively. The reciprocity of *i* is the fraction of all neighbors that are both upstream and downstream, 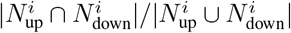 (Fig. 5g). Reciprocity is bounded between 0 (fully feedforward) and 1 (fully recurrent).

## Data Availability

Raw connectome data analyzed in this study are previously published and are available as deposited at Zenodo (https://doi.org/10.5281/zenodo.23427196).

## Code Availability

The code used to perform the analyses in this paper is openly available at github.com/jbetancourt015/robust_communication.

## Supplementary Information

Supplementary text and figures accompany this paper.

## Acknowledgements

We thank Sekhar Tatikonda, Sanjana Taneja and Bryce Lowden for very informative discussions. J.M.B. acknowledges support from the Physics, Engineering, and Biology Program at Yale University. C.W.L. acknowledges support from the National Institutes of Health (NIH/NIGMS R35GM160188) and the Wu Tsai Institute at Yale University.

## Author Contributions

J.M.B., V.V., and C.W.L. conceived the project; J.M.B. and C.W.L. designed the models; J.M.B. performed the analysis with input from C.W.L. and D.A.C.; and J.M.B. and C.W.L. wrote the paper with input from D.A.C. and V.V.

## Supplementary Information

### 1. Introduction

In this Supplementary Information, we provide additional analyses, statistics and robustness checks to support the results in the main text. Throughout, we use E_*u*_(·) to denote the expectation value with respect to the random variable *u*; angle brackets ⟨·⟩ instead denote empirical averages over a neuron’s realized incoming connection weights. In Sec. 2, we provide an alternative derivation of the robustness result that makes use of the geometric intuition of classification planes in the space of inputs. In Sec. 3, we extend our results to sparse inputs and outputs, showing that the input weights still only enter the error rate through the robustness of the neuron. In Sec. 4, we give conditions under which robustness characterizes the error rate when inputs are correlated. In Sec. 5, we show that, under a more general class of noise structures, the error rate still depends on the input weights only via (a modified version of) robustness. In Sec. 6, we extend our analysis to continuous outputs, and show that the advantages of heavy-tailed connection strengths arise from the nonlinearity of neuronal responses. In Sec. 7, we demonstrate that our error rate prediction remains accurate under alternative definitions of connection strengths. In Sec. 8, we show that high robustness arises from the distributional structure of incoming connection strengths. In Sec. 9, we show how different choices of shuffling procedures affect robustness when connections are thresholded. In Sec. 10, we measure the fraction of synapses that get assigned the wrong brain region when connections and neurons are assigned to regions, and show that the majority of our coarse-grainings have a small fraction of misclassified synapses.

### 2. A geometric visualization of robustness

Our analysis of the error rate relied on analyzing the distribution of the local field *z* and its perturbations 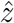. We can build a complementary intuition by analyzing the effects of perturbations in input space.

Before we begin our analysis, we note that our derivation of the analytic prediction of the error rate does not rely on the inputs being binary, only that they have zero mean. Therefore, our prediction for the error rate also holds for centered continuous inputs. This allows us to establish a clean benchmark for our geometric analysis: Gaussian inputs. Once we have built intuition for this benchmark, we can apply it to our binary input framework.

We consider inputs drawn from a multivariate Gaussian:

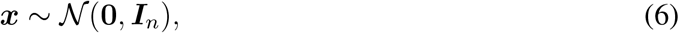

where ***I***_*n*_ is the identity matrix of dimension *n*. Given this distribution of inputs, we can ask: what is the probability of an error for weights ***w*** and a given set of perturbed weights 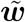? The answer is that the error is proportional to the angle *θ* between the perturbed and unperturbed weights (Eq. 4.12 in Ref. 65):

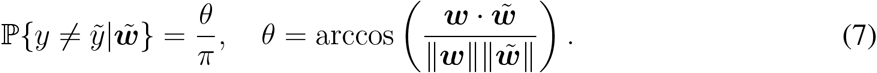

This is easy to visualize in the case of two inputs. Consider first only the inputs at a distance *R*. Given the rotational symmetry of the Gaussian, the distribution of inputs at a given distance is uniform. Therefore, the probability of an error given that the input is at a distance *R* is the fraction of the circle on which errors occur, which is 2*θ/*2*π* = *θ/π* (Fig. S1a). For *n* dimensions, we can compute this probability by segmenting space into planes parallel to the plane formed by ***w*** and 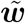. Conditional on the input lying on one of these planes, we can use an argument analogous to the two-dimensional one to show that the error probability is *θ/π*. Since this is true for any plane, it is true for all space, and we obtain the general result in Eq. (7).

We now have a geometric interpretation for what is driving the errors for a fixed perturbation. How do we extend this interpretation to the error rate? To obtain the error rate, we have to average over all possible 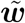:

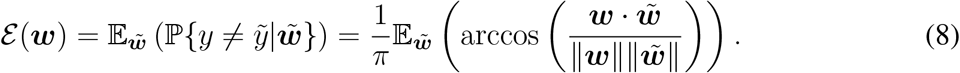

Since we are interested in the large-*n* behavior of the error rate, it is useful to re-write the expression above in terms of averages rather than sums, since we can expect these not to scale with *n*.

**Fig. S1.**
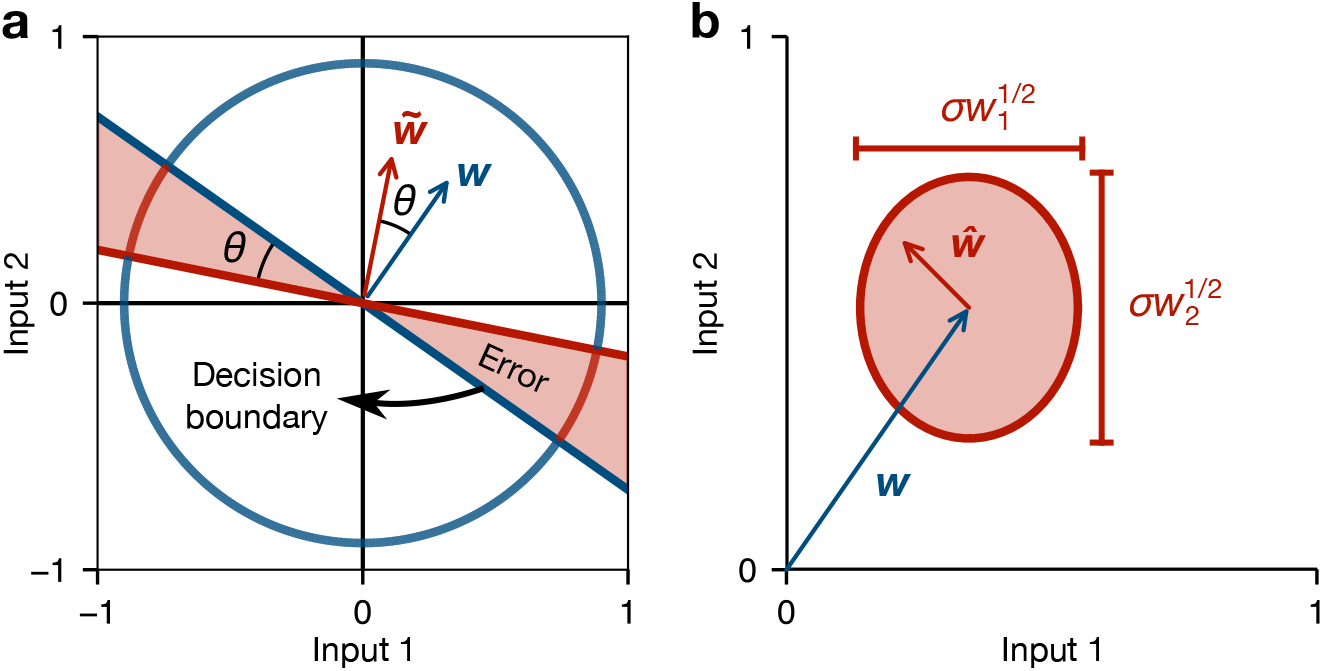
Geometric visualization of communication errors. **a**, Perturbations in the weights ***w*** shift the decision boundary of the neuron. When inputs are rotationally-symmetric, the probability of an error is proportional to the angular shift of the decision boundary. **b**, Input weights ***w*** define the distribution of weight perturbations ***ŵ***, which determines the distribution of the angle *θ* between ***w*** and the perturbed weights 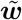. The shaded red area shows a level set of the distribution of perturbed weights.

Letting 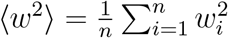, we can write cos(*θ*) as

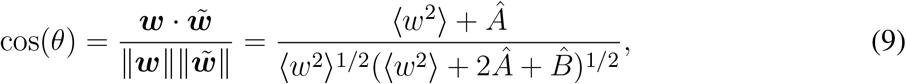

where 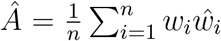 and 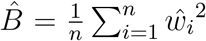. For large *n*, we expect the quantities *Â* and 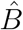 to concentrate around their average values (assuming ⟨*w*⟩ and ⟨*w*^2^⟩ converge):

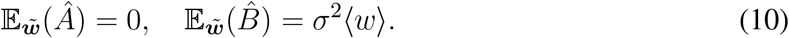

Indeed, all central moments of *Â* and 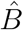 are at most *O*(1*/n*). Therefore, expanding *θ* in terms of *Â* and 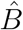 around their averages yields a large-*n* expansion of the error rate:

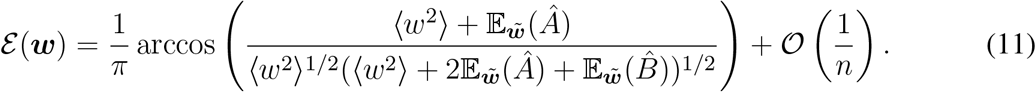

Using the expectations in Eq. (10) and dropping higher-order terms, we obtain our analytical prediction of the error rate:

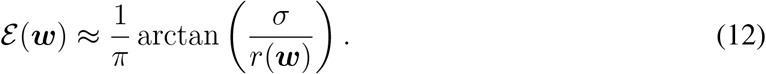

The analysis above gives an alternate intuition for what drives errors. Perturbations generate a distribution of weights centered around the unperturbed weights ***w*** (Fig. S1b). The perturbations ***ŵ*** are, in expectation, orthogonal to the original vector 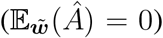. Thus, what determines the magnitude of *θ* is mostly the magnitude of perturbations in the orthogonal direction 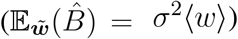. These intuitions about errors being related to the angles between decision boundaries apply roughly to our binary input setup. While the results for fixed perturbations are specific to rotationally-invariant distributions (like the Gaussian), we get the same quantitative behavior once we average over perturbations.

### 3. Sparse inputs and outputs

Our prediction of the error rate assumes that there is no bias in the local field and that inputs are zero-mean. This implies that the unperturbed neuron fires half the time, since the symmetry implies P{*z >* 0} = 1*/*2. We can use our framework to explore how the error rate changes when we allow inputs and outputs to be sparse, and how the distribution of weights determines this behavior.

To relax the assumption of symmetric inputs, we now allow inputs to be drawn independently from a distribution with mean *µ*_*x*_ and variance 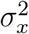. For example, this encompasses our original binary input setup with different probabilities of firing and not firing. We still assume that inputs are independently drawn. If the probability of staying silent is higher than the probability of firing, then *µ*_*x*_ *<* 0. We also relax the assumption that there is no bias in the local field, which affects the firing probability of the postsynaptic neuron. We introduce a bias *b* to the local field:

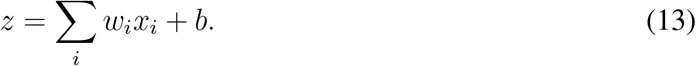

Decreasing the bias leads to a decrease in the firing rate of the neuron.

Allowing sparse inputs and outputs, we again make progress by studying the *n* → ∞ limit. However, in this limit we must be careful about the scaling of *b* and *µ*_*x*_. Rather than keeping *µ*_*x*_ and *b* fixed, we instead keep the probability that the neuron fires, which we call *p*_*f*_, constant as the number of inputs *n* grows.

We begin our asymptotic analysis by studying the distribution of the unperturbed local field. It is useful to define the centered local field *z*_*c*_ = ∑*i w*_*i*_(*x*_*i*_ − *µ*_*x*_) and characterize its asymptotic behavior. As we let *n* grow, we assume weights are drawn independently and that *σ*_*x*_ is held constant. The results, however, also hold if *σ*_*x*_ is allowed to vary but converges to an *O*(1) value. Under these assumptions, we can apply the central limit theorem to conclude that the centered local field is asymptotically Gaussian:

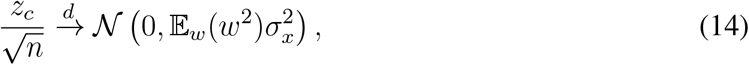

where 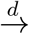 represents convergence in distribution. Note that this result holds regardless of the asymptotic behavior of *µ*_*x*_ and *b*.

Knowing the behavior of *z*_*c*_ allows us to study the distribution of *z*, even for finite *n*. For a fixed ***w***, note that the expected value of *z*, which we call *µ*_*z*_, is *µ*_*z*_ = *nµ*_*x*_⟨*w*⟩ + *b*. Therefore, we can decompose the local field as

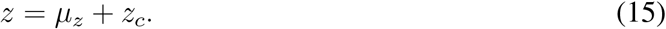

This decomposition allows us to characterize the firing rate *p*_*f*_ . The probability of firing is

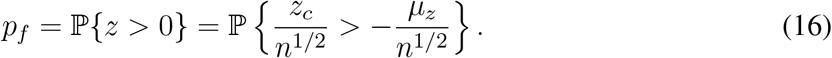

We can use the asymptotic behavior of *z*_*c*_*/n*^1*/*2^ to approximate the firing rate (Fig. S2a). For a given ***w***, we use E_*w*_(*w*^2^) ≈ ⟨*w*^2^⟩ to obtain

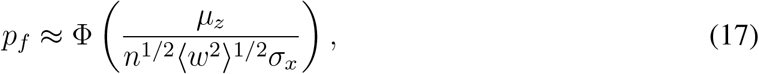

where Φ is the CDF of the standard normal distribution. Note that to maintain a firing rate *p*_*f*_ ≠ 1*/*2 that does not approach 0 or 1, we must have *µ*_*z*_ ∝ *n*^1*/*2^. Thus, a sparse, but finite, firing rate can be achieved with *µ*_*x*_ ∝ *n*^−1*/*2^ or *b* ∝ *n*^1*/*2^.

Having characterized the behavior of the unperturbed local field in terms of the firing rate, we can now study the effect of perturbations in sparse neurons. It is useful to work with the centered version of the perturbed local field: 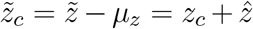. We can use the same convergence result as in the main text for the joint distribution of 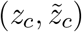:

**Fig. S2.**
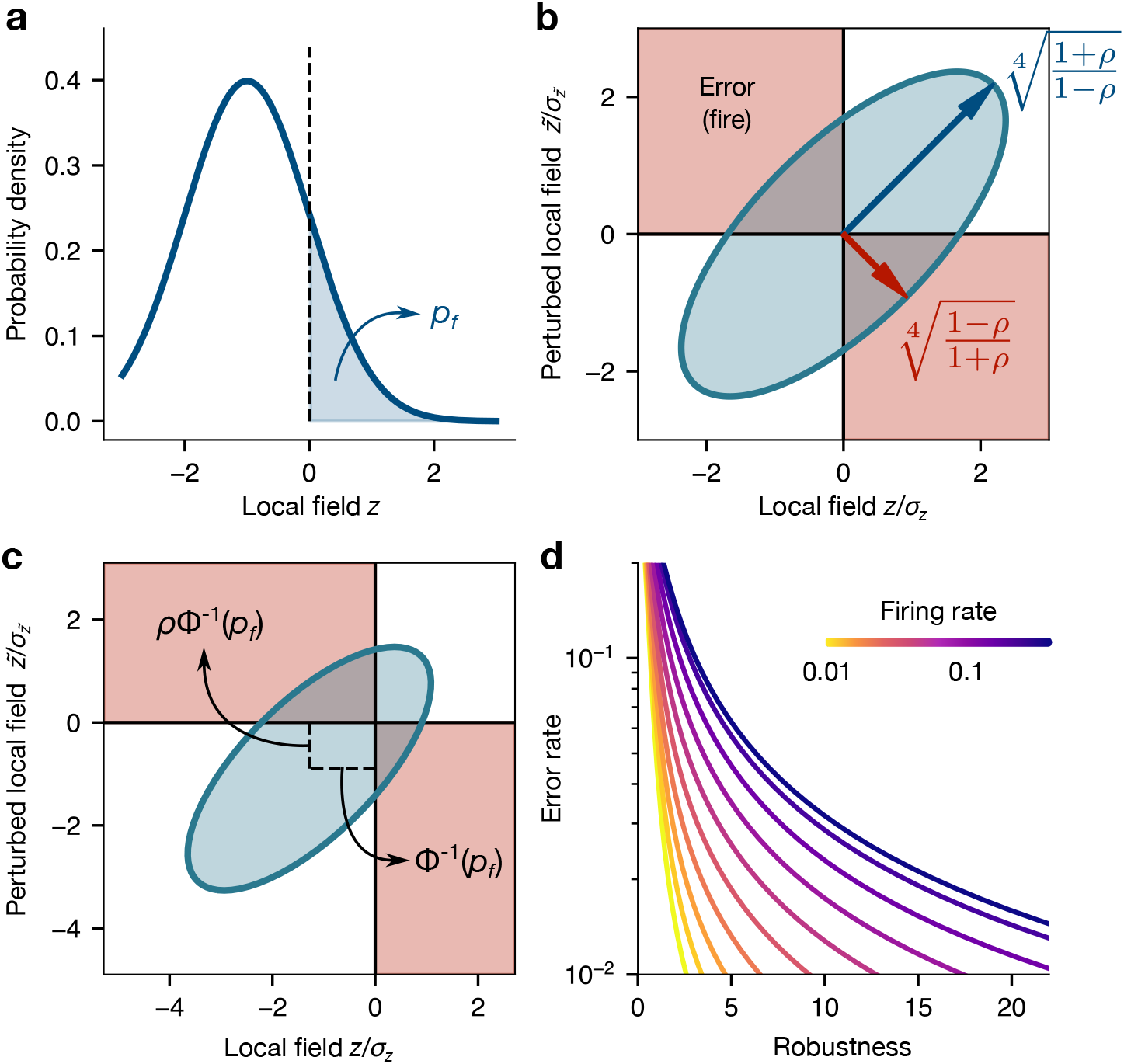
Geometric visualization of errors when inputs and outputs are sparse. **a**, Under non-symmetric inputs or non-zero bias, the local field distribution can be characterized asymptotically by the firing probability *p*_*f*_ = P{*z >* 0}. **b**, Errors occur in the second and fourth quadrants in the space of local fields and perturbed local fields. For symmetric inputs and zero bias, the joint distribution of 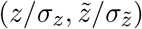 is fully characterized by the correlation *ρ* between *z* and 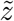. The shaded blue region represents a level set of the distribution. Arrows show the principal axes of the elliptic level set. **c**, When the unperturbed firing probability is *p*_*f*_ ≠ 1*/*2, the distribution of 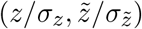 is shifted away from the origin. **d**, Error rate vs robustness for perturbation strength *σ* = 1 at varying firing rates *p*_*f*_ .

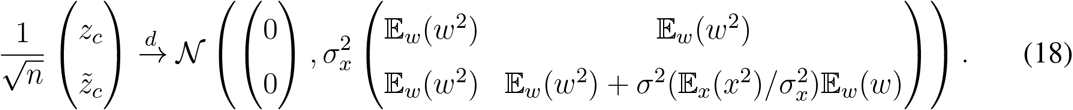

For our analysis, in which the local field has non-zero mean, it is useful to state this result in terms of the normalized version of the variables:

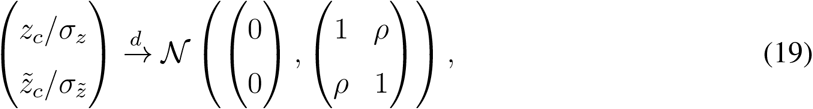

where the variances are

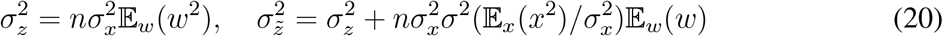

and *ρ* is the correlation between *z*_*c*_ and 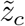:

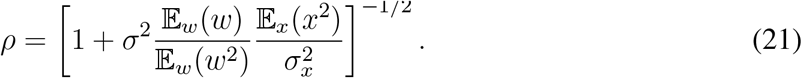

In 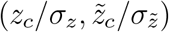 space, the correlation *ρ* sets the shape of the asymptotic joint distribution. In this space, the level sets of the joint distribution are ellipses whose major axis lies on the *y* = *x* line. The correlation sets how much the ellipse is stretched along this axis, with a higher correlation leading to a more elongated ellipse. For *p*_*f*_ = 1*/*2 this elongation sets an intuitive picture for understanding the asymptotic role of the correlation in the error rate (Fig. S2b).

With our characterization of the asymptotic behavior of *z*_*c*_ and 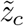, we can analyze the distribution of 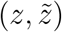 to obtain the error rate. Note that we can (asymptotically) write the mean of *z* in terms of the firing probability *p*_*f*_ :

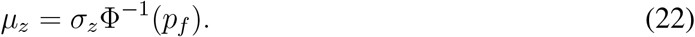

Using this result, the means of our normalized local fields are

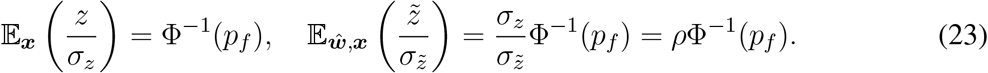

With this result, we see that the joint distribution of 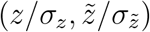 is fully characterized by the correlation *ρ* and the firing rate *p*_*f*_ (Fig. S2c). Therefore, the asymptotic error rate is also a function of only *ρ* and *p*_*f*_ .

To compute the error rate E(*ρ*; *p*_*f*_ ), we decompose the error probabilities as

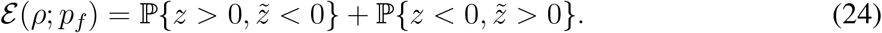

Note that

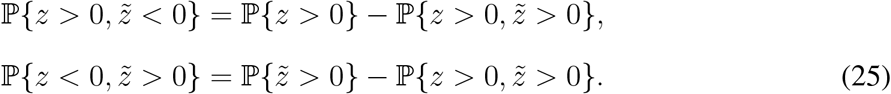

Therefore, the error rate is

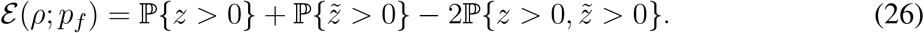

Equivalently, in terms of normalized variables,

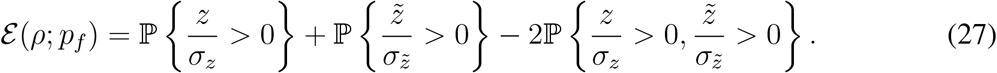

Using our characterization of the joint distribution of the normalized local fields, these probabilities can be written explicitly:

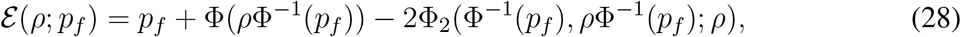

where Φ_2_ is the bivariate standard normal CDF. As mentioned above, this error rate depends only on *ρ* and *p*_*f*_ . For finite *n*, we can approximate the correlation using robustness:

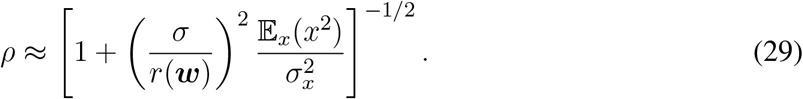

Thus, we have an analytical prediction of the error rate for sparsely-firing neurons that depends only on robustness and the firing rate. Specifically, the error rate is decreasing in robustness (as before) and increasing in the firing rate (Fig. S2d).

### 4. Correlated inputs

In this Section, we study the asymptotic behavior of the error rate when inputs are allowed to be correlated. We restrict ourselves to the case of *x*_*i*_ ∈ {−1, +1} and *b* = 0. We also assume that E_*x*_(*x*_*i*_) = 0, which implies that Var(*x*_*i*_) = 1. For a fixed weight vector ***w***, the covariance matrix of the local field *z* and perturbed local field 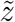 conditional on ***w*** is

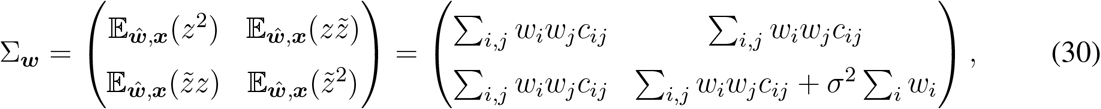

where *c*_*ij*_ = Cov(*x*_*i*_, *x*_*j*_). If we now assume that the weights ***w*** are drawn iid from a distribution *P* (*w*), and independently from the inputs, the unconditional covariance matrix is

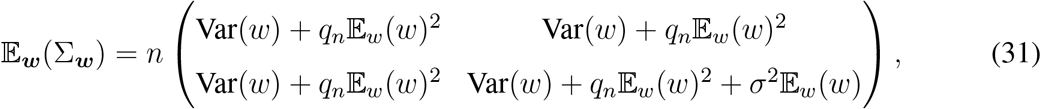

where 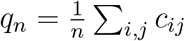. Thus, the correlation between *z* and 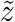 at a finite number of inputs is

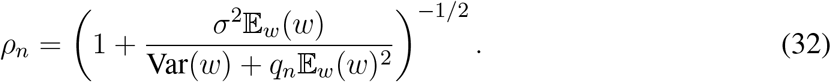

We can write this correlation as 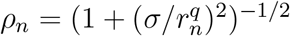, where we now use a modified version of robustness:

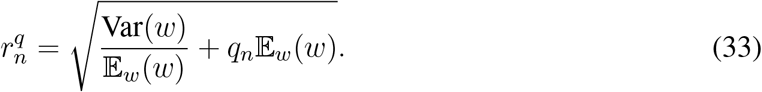

When inputs are uncorrelated, we have *q*_*n*_ = 1 and we recover our previous result. Robustness is still monotonically increasing with variance, and is monotonically increasing in the off-diagonal correlations between inputs.

While we can intuitively characterize the correlation between *z* and 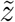 when inputs are correlated, the independence condition needed to apply the central limit theorem is violated. To extend our covariance result to a prediction about the error rate, we need to establish weaker conditions for convergence to Gaussianity.

We first establish conditions under which 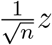 converges to a Gaussian as *n* → ∞. We split the local field into two contributions:

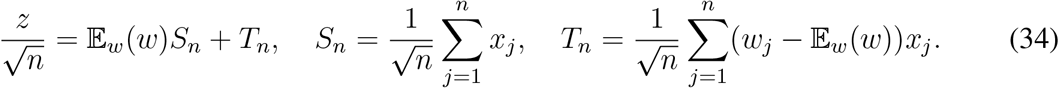

This decomposition allows us to break down our analysis into characterizing the asymptotic properties of two simpler objects. Intuitively, we can show that *T*_*n*_ converges to a Gaussian regardless of the distribution of ***x***. The difficulty is in obtaining sufficient conditions for *S*_*n*_ to also converge to a Gaussian.

To prove convergence we use a characteristic function approach. The characteristic function of the distribution of a single weight is

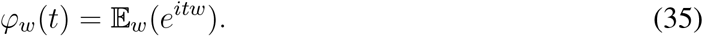

It is useful to characterize the cumulant generating function log(*Φ*_*w*_(*t*)). Expanding this function to second order yields

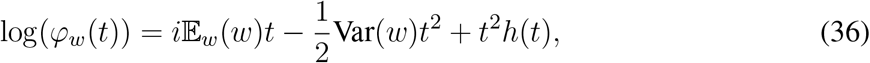

with *h* → 0 as *t* → 0.

Before studying the full characteristic function of 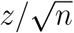, we compute the expectation of 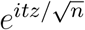 conditioned on an input realization ***x***. Since the weights are independent and the *x*_*j*_ are constant under the conditioning, the expectation factors as

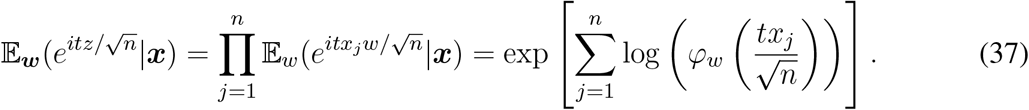

Using our expansion of the cumulant generating function, we obtain

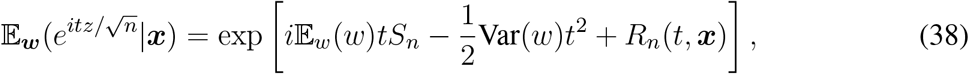

where the aggregate residual is

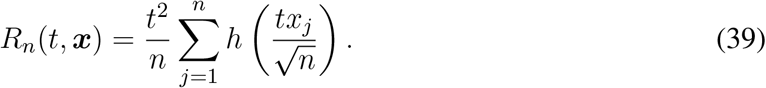

Note that we can bound the size of this residual by

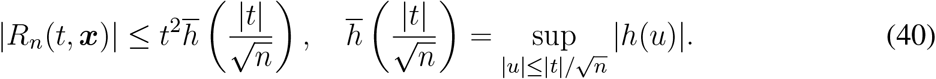

Importantly, note that the residual vanishes as *n* → ∞ for all realizations ***x*** at any fixed *t*. Using our bound on the residual, we obtain the following for all inputs ***x***:

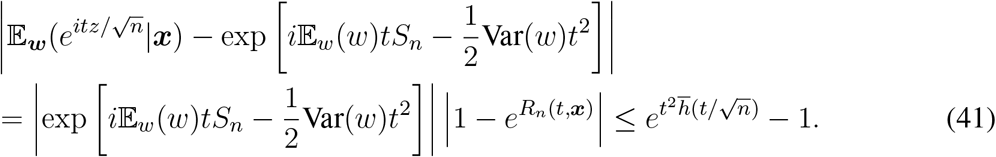

Since the distance bound above is valid for all ***x***, it is also valid for the averages with respect to ***x***:

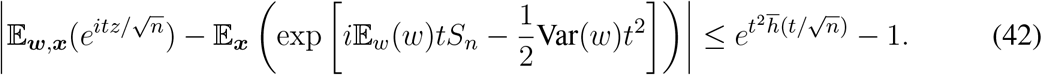

Note that 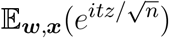 is the characteristic function of 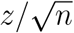 and 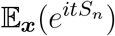 is the characteristic function of *S*_*n*_. Therefore, our bound reads

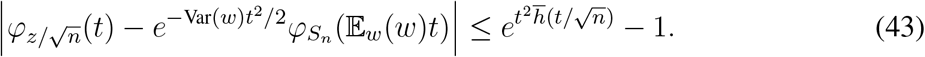

Up to this point we have made no assumptions on the joint distribution of ***x***, other than *x*_*i*_ ∈ {−1, +1} and E_*x*_(*x*_*i*_) = 0. To establish convergence to Gaussianity, we need the characteristic function to converge to asymptotically have the form 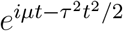. This means that our modified central limit theorem requires that *S*_*n*_ converges to a Gaussian. We know E_***x***_(*S*_*n*_) = 0 and

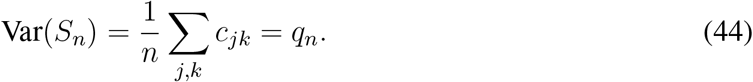

Therefore, we can guarantee *S*_*n*_ converges to a Gaussian if *q*_*n*_ → *q* for some fixed *q* and

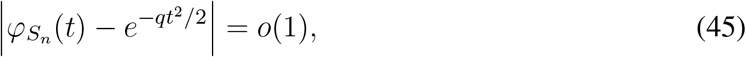

where *o*(1) means the value goes to 0 as *n* → ∞. Under these conditions, we can use our previous distance bound to conclude that the characteristic function of 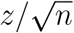 satisfies

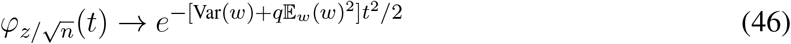

for all *t*. The convergence of the characteristic function implies convergence in distribution, meaning that

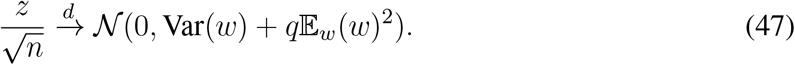

What conditions are sufficient for *S*_*n*_ to converge to a Gaussian? A basic necessary condition is that *q*_*n*_ needs to remain bounded, which means that the average correlation 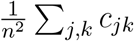 must go to 0. However, this is not a sufficient condition. An intuitive sufficient condition can be formulated in terms of dependency graphs. In a dependency graph of random variables, any two disconnected sets of vertices correspond to mutually independent families of variables. Note that dependency graph need not be unique. Defining *D*_*n*_ as the maximum degree of a dependency graph of ***x***, a sufficient condition for Gaussian convergence is that *D*_*n*_ = *o*(*n*^1*/*4^) (Theorem 4.3 in Ref. 66). Intuitively, this condition guarantees that there are no “global” couplings that can collectively perturb the distribution of *z*. For example, if the inputs were perfectly correlated, *z* would be composed of two Diracs. There also exist conditions on the (conditional) correlations between variables (Theorem 1 in Ref. 67), which are not as strict as independence but are significantly more complex to evaluate. These conditions encode the intuition that learning about one variable cannot reveal too much information about the rest of the variables.

We now return to the distribution of 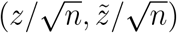 and analyze its convergence. To extend our result to the multivariate case, we use the Cramér-Wold theorem,^68^ which formulates sufficient conditions for multivariate convergence starting from single-variable convergence. For our application, to prove that 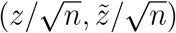 converges to a Gaussian with variances 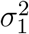 and 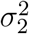 and covariance *C*, it is sufficient to prove that the following holds for all *α, β* ∈ ℝ:

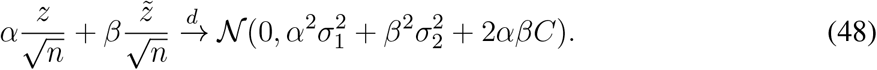

We can decompose the relevant linear combination as

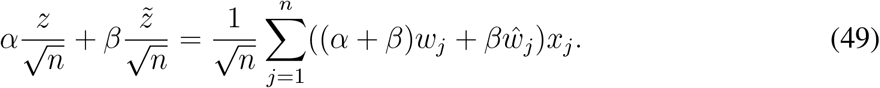

Note that the random variables (*α* + *β*)*w*_*j*_ + *βŵ*_*j*_ are iid, so we can reproduce the proof for convergence in distribution of *z* by using the replacement *w* → (*α* + *β*)*w* + *βŵ*. We need the following statistics:

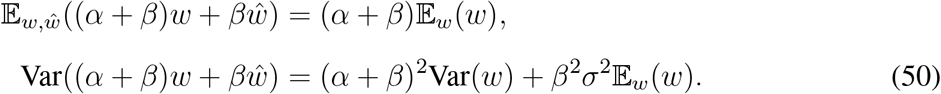

Therefore, under the same assumptions that give convergence of *z*, we obtain

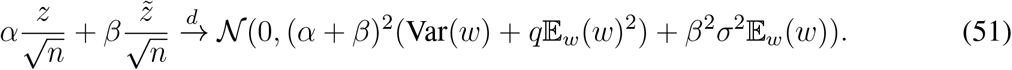

We conclude that the joint distribution satisfies the following convergence result:

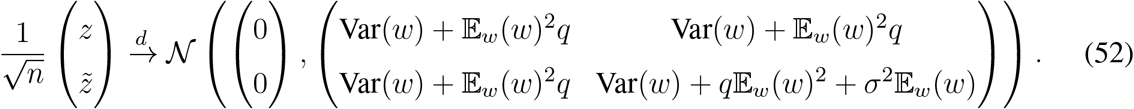

With this result, we recover our error rate with a modified robustness metric:

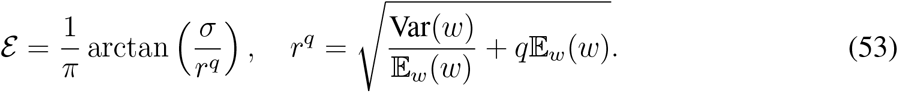

### 5. Other noise structures

In the main text, we focused on errors induced by additive noise at the synapse level. However, our framework allows for a more general weight perturbations. We still assume that the perturbations *ŵ*_*i*_ satisfy E_*ŵ*_(*ŵ*_*i*_) = 0 and are independently drawn. However, we now assume that their variance has a general structure,

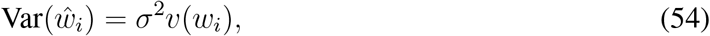

where *v* is an arbitrary function of the weights. The only condition we impose on *v* is that it does not make the variance of 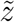 grow unbounded. We maintain the noise strength parameter *σ* for expositional clarity. The joint distribution of 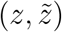 now satisfies

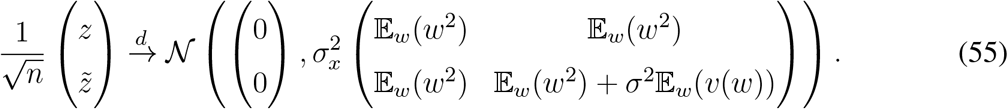

The computation of the error rate then follows as in the main text. This implies that, for fixed ***w***, we can make a prediction of the error rate *ε*_*v*_(***w***) associated to this noise structure:

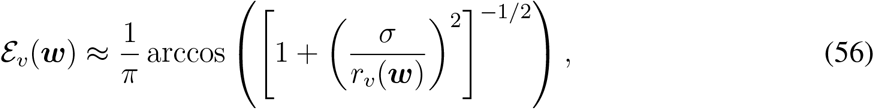

where robustness now has the modified form

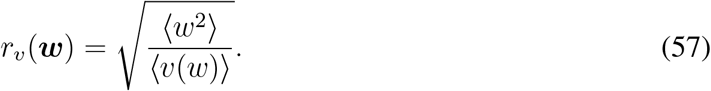

Therefore, the simplicity of our robustness prediction is not tied to our particular choice of noise structure; the error rate still depends on the weight only via a simple statistic. The noise structure only determines the specific form that robustness takes.

Consider noise of the form

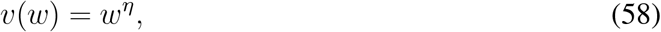

which produces robustness,

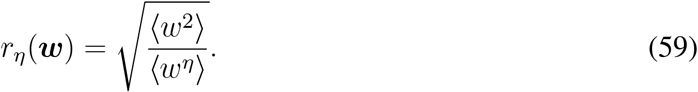

Note that the additive noise we consider in the main text corresponds to *η* = 1. This characterization allows for two additional important classes of noise. For *η* = 0, we obtain perturbations that have the same strength regardless of the weight. This constant noise could arise from processes that affect communication at the connection level, such as fluctuations at the level of entire presynaptic neurons. In this scenario, robustness monotonically increases with both the average connection strength ⟨*w*⟩ and variance Var(*w*). Another case of interest is *η* = 2, which corresponds to multiplicative noise. Multiplicative noise arises when there is a common noise source shared by all synapses, which is then amplified by each additional synapse. In this scenario, the robustness becomes independent of the connection strengths.

We can study which distributions of weights give a higher or lower robustness at a given *η*. For this, we fix the average weight ⟨*w*⟩ and ask which distributions have the highest or lowest robustness. For *η* = 1, we showed that the weight distribution with the lowest robustness is the uniform distribution, in which *w*_*i*_ = ⟨*w*⟩. In fact, this is true for all sub-multiplicative noise, *η* ∈ [0, 2]. To see this, note that by Jensen’s inequality,

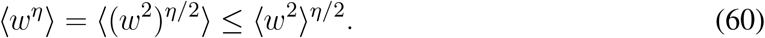

Using this inequality and ⟨*w*^2^⟩ ≥ ⟨*w*⟩^2^ (also by Jensen’s inequality), we see that robustness satisfies

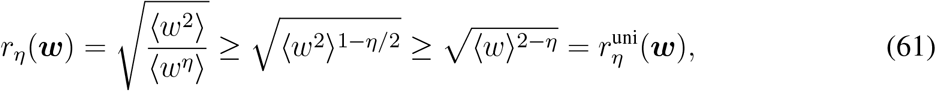

where 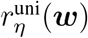 is the robustness value of the uniform weight vector. We can also characterize the weights that maximize robustness. Assuming the weights are non-negative, the largest possible value of a given *w*_*i*_ is *n*⟨*w*⟩. With this, we can obtain a lower bound for ⟨*w*^*η*^⟩:

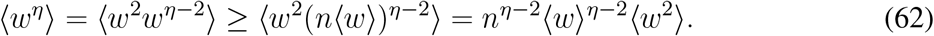

Thus, we obtain an upper bound for robustness:

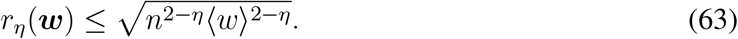

This bound can be achieved by the maximally-sparse weight vector: ***w*** = (*n*⟨*w*⟩, 0, …, 0) or any of its permutations. Calling the robustness of the single-entry weight vector 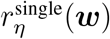, we see that, in general,

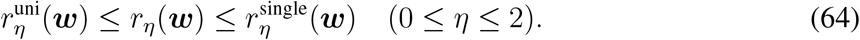

For larger values of *η*, the relevant inequalities flip, giving the reverse order:

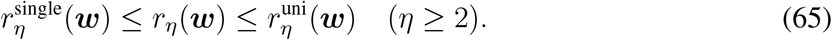

Thus, for sub-multiplicative noise *η* ∈ [0, 2], high robustness requires the distribution of weights to be heavy-tailed. This is because distributions with heavier tails create wider distributions of local fields without producing a comparable effect in the distribution of the local field perturbation 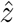 (Fig. 2c in the main text). In contrast, when noise is super-multiplicative (*η >* 2), widening the distribution of the local field requires a super-linear widening of the local field perturbation, inducing more errors. Therefore, in this case it is beneficial to have the lowest possible spread in connection strengths, corresponding to the uniform distribution.

### 6. Communication with continuous output

Our main analysis focused on binary inputs, motivated by spiking neurons. However, we can extend our framework to analyze how perturbations induce errors in graded responses. As a simple example, consider neurons with a sigmoidal response to the local field. Motivated by our symmetric binary convention (+1 for firing and −1 for silence), we take the output to be

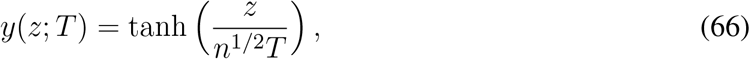

where *T* is a temperature parameter that modulates the size of the region in *z* space in which intermediate values of *y* are possible. This is because for 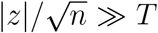 the outputs are essentially binary. The 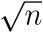 scaling is introduced to avoid trivial behaviors in the model in the large *n* limit.

To quantitatively study the effects of perturbations in a continuous output setting, we need to generalize the concept of error rate used for binary neurons. We study a simple quadratic loss, where the pointwise loss obtained from a particular realization of an unperturbed output *y* and a perturbed one 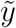 is

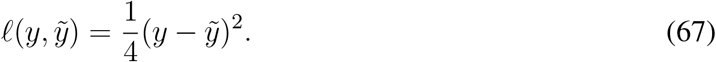

While we focus on this particular form of loss, our analysis would remain qualitatively the same for any smooth loss function. Note that when *T* = 0, this quadratic loss coincides with the misclassification loss 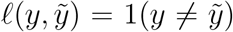, where 1(·) is the indicator function. The performance of a neuron is quantified by its average loss (or simply loss)

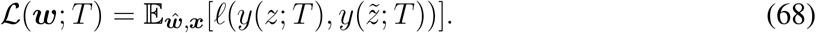

As the temperature parameter approaches 0 the loss simply becomes our original error rate:

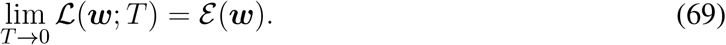

The sigmoidal activation naturally gives rise to two regimes. As mentioned above, 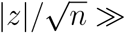 corresponds to a saturated regime, in which the output is essentially binary, and small changes in the local field do not affect the output. By contrast, when 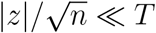 the output is approximately a linear function of the local field, meaning that small changes in the local field have proportional effects on the output. This behavior translates to a separation of 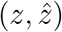 space into regions where outputs are approximately linear or binary. For a given ***w***, the qualitative behavior of the loss depends on how much the distribution of 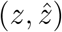 overlaps with the linear region (Fig. S3). For large temperatures, the bulk of this distribution is in the linear region, so the loss takes the form

**Fig. S3.**
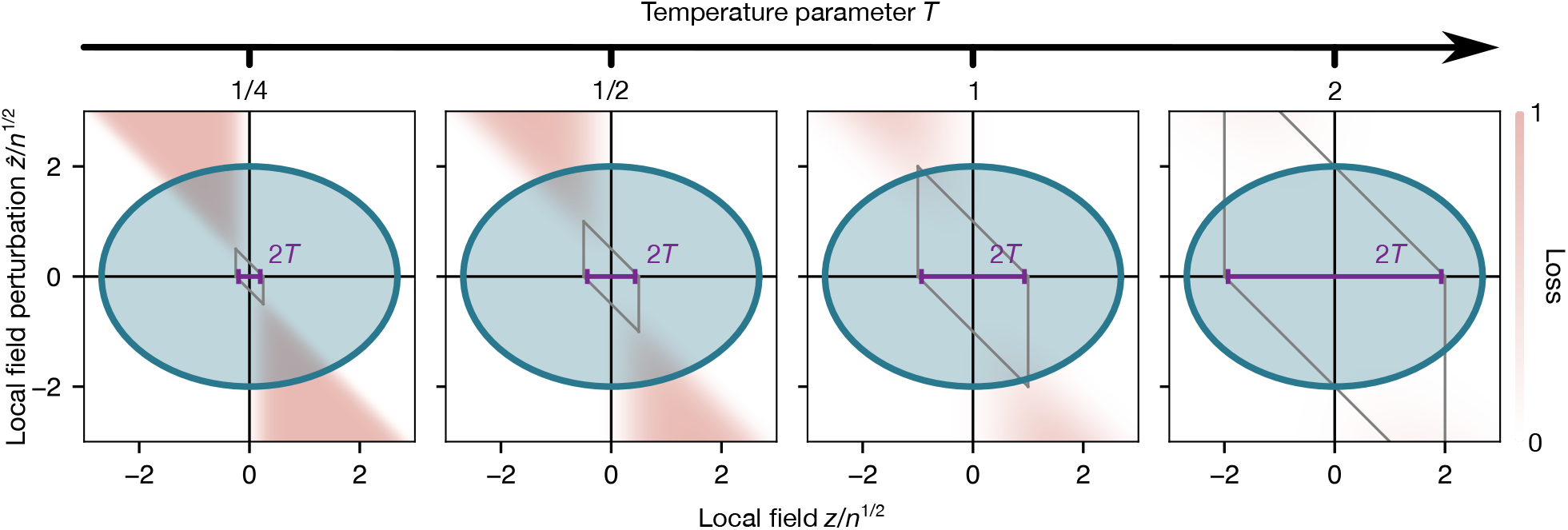
Visualization of loss under continuous outputs. The joint distribution of the local field and the local field perturbation (level set shown in blue) determine the loss. The temperature parameter *T* sets the size of the linear region (gray parallelogram). For very low temperatures, the linear region is much smaller than the scale of the distribution, and the loss behaves as the binary loss in the main text. For very large temperatures, the bulk of the distribution is in the linear region, and the loss becomes proportional to 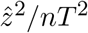.

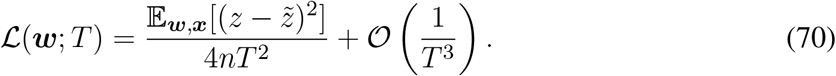

Taking the expectation, we get

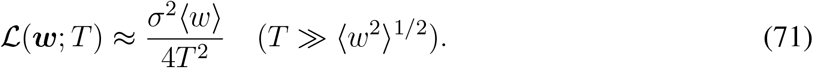

This result is in striking contrast to the loss at small *T* . In this linear regime, there is no benefit to having a large variance in weights: all weights ***w*** with the same ⟨*w*⟩ have approximately the same error. This is because widening the distribution of local fields no longer has the sharp effect of changing outcomes from an “error” region to a “correct” region. Instead, the linearity of the output means that for a fixed perturbation, changing the local field does not affect the loss. As the temperature parameter decreases, nonlinear effects become important and the advantages of having a wide distribution of weights become stronger. This result shows that the advantages of heavy-tailed distributions of connection strength are inherently tied to the nonlinearity of neuron responses.

**Fig. S4.**
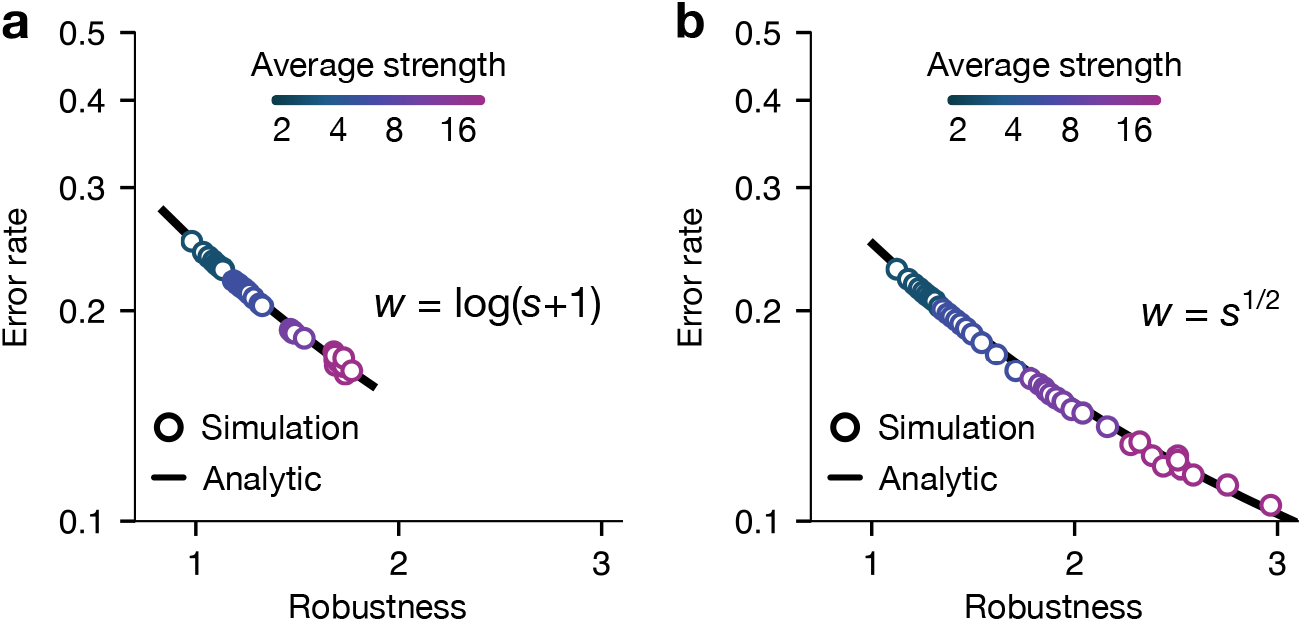
Error rates are determined by robustness in alternative definitions of connection strengths. **a**, Error rates versus robustness for neurons in the fruit fly brain^7,8^ with at least ten incoming connections using connection strengths *w* = log(*s* + 1). Neurons were grouped into the same groups as Fig. 3 of the main text. The median simulated error rate and median robustness of each group are shown as scatter points. The solid line represents our analytical prediction. **b**, Error rates versus robustness using connection strengths *w* = *s*^1*/*2^. For both scenarios, the perturbation strength is *σ* = 1.

### 7. Alternative definitions of connection strengths

In the main text, we used two different definitions of connection strength: synapse count for invertebrates and contact area for mouse retina. However, our framework allows for more general definitions of connection strengths. To illustrate this generality, we study errors for connection strengths of the form *w* = *f* (*s*), where *s* is the number of synapses in a connection. Specifically, we consider *w* = log(*s* + 1) and 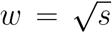. We find that our prediction for the error rate is in excellent agreement with simulations and that the specific choice of *f* only changes the distribution of robustness, but not the accuracy of our analytic prediction (Fig. S4). In simulations with *w* = log(*s* + 1), the predicted error rate is within 0.004 of the simulated one on average, compared to an average simulated error rate of 0.221. For 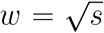, predicted error rate is within 0.004 of the simulated one on average, compared to an average simulated error rate of 0.197.

**Fig. S5.**
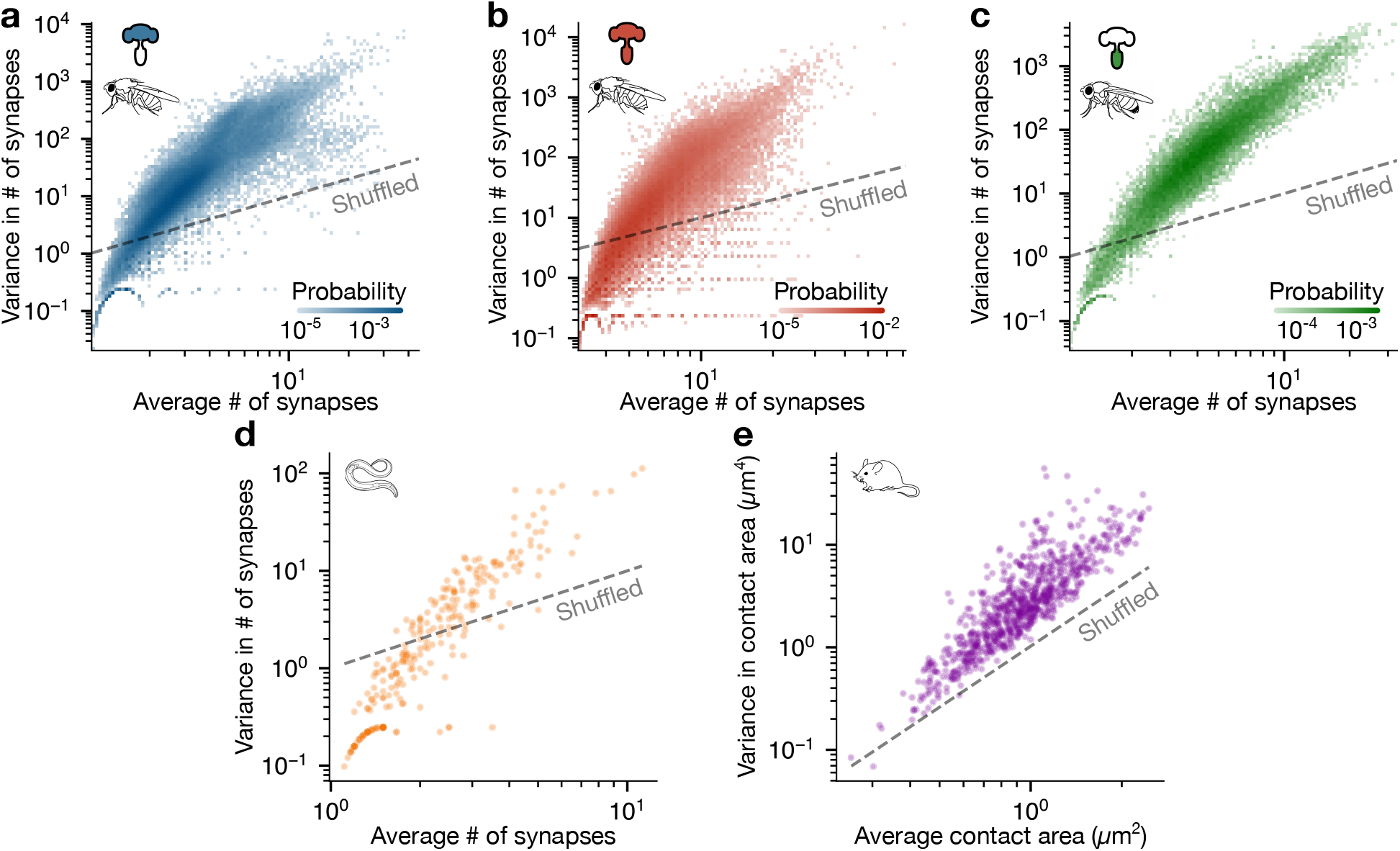
Connection strength statistics across species. **a**-**d**, Average number of synapses and variance in the number of synapses of the incoming connections of neurons in the female fruit fly brain (**a**),^7,8^ the combined brain and nerve cord of a *Drosophila* female (**b**),^9^ the *Drosophila* male nerve cord (**c**),^10^ and the nematode *C. elegans* (**d**).^3,4^ **e**, Average contact area and variance in the contact area of incoming connections of neurons in the inner plexiform layer of the mouse retina.^5^ Dashed lines represent the variance-mean relationship after connections are shuffled (Var(*w*) = ⟨*w*⟩ for invertebrates, Var(*w*) = ⟨*w*⟩^2^(*k*_in_ − 1)*/*(*k*_in_ + 1) ≈ ⟨*w*⟩^2^ for the mouse retina).

We expect our error rate prediction to remain accurate for other choices of *f* . For our results to remain valid, the conditions for the convergence of the 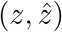 distribution must still hold. The central concern is that the distribution of *w* becomes too wide and large weights dominate the output. Formalizing this intuition, we can show that any *f* that reduces the spread in the weights guarantees convergence. For example, this is possible if *f* satisfies |*f* ^′^(*s*)| ≤ 1, as in the cases studied above.

### 8. Distribution of connection strength statistics

As shown in the main text, real neurons are consistently more robust than neurons with random synaptic connectivity. The robustness of a neuron is determined by the statistics of its incoming connection strengths. Thus, by analyzing the distribution of incoming statistics across neurons, we can obtain a more detailed picture of what drives their high robustness. We see that the joint distribution of means and variances in incoming connection strengths is consistently concentrated along a one-dimensional relationship across organisms, such that a high average connection strength is accompanied by a high variance (Fig. S5). This concentrated distribution of statistics is consistently above the Var(*w*) = ⟨*w*⟩ benchmark, which corresponds to shuffling units of connection strength among connections. This concentration above the shuffled benchmark results in high robustness for most neurons.

### 9. Shuffling synapses in thresholded connectomes

In the connectome of the combined brain and nerve cord of a *Drosophila* female,^9^ connections are thresholded, such that only connections with at least three synapses are reported. Thresholding introduces additional choices on how to construct null neurons for our analysis in Fig. 4 of the main text. There, we construct shuffled neurons by first placing three synapses into each incoming connection and then placing synapses uniformly at random until the total number of incoming synapses matches that of the real neuron. In this Section, we propose two alternative shuffling schemes and compare the robustness of the resulting neurons against the robustness of the real neurons.

In the first alternative scheme, synapses are shuffled in groups. Intuitively, we can interpret the threshold as imposing a resolution on synapse counts. We can account for this imposed resolution by placing synapses in groups of three uniformly at random, until the number of synapses matches the total number of the original neuron. The second alternative is to simply use the same scheme as the other invertebrate connectomes: assigning synapses to incoming connections uniformly at random until the total synaptic count matches the original neuron.

The top row of Fig. S6 shows the robustness of neurons before and after shuffling synapses using the above schemes. We see that the fraction of neurons whose robustness is higher than their shuffled counterparts is lower than in Fig. 4 of the main text. This is because these alternative shuffling procedures can leave connections with zero synapses, which allows the distribution of connection strengths to have a higher variance and, hence, higher robustness. Intuitively, this is more important in neurons that have connections with low synaptic counts. This explains why only low-robustness neurons increase in robustness when shuffled.

**Fig. S6.**
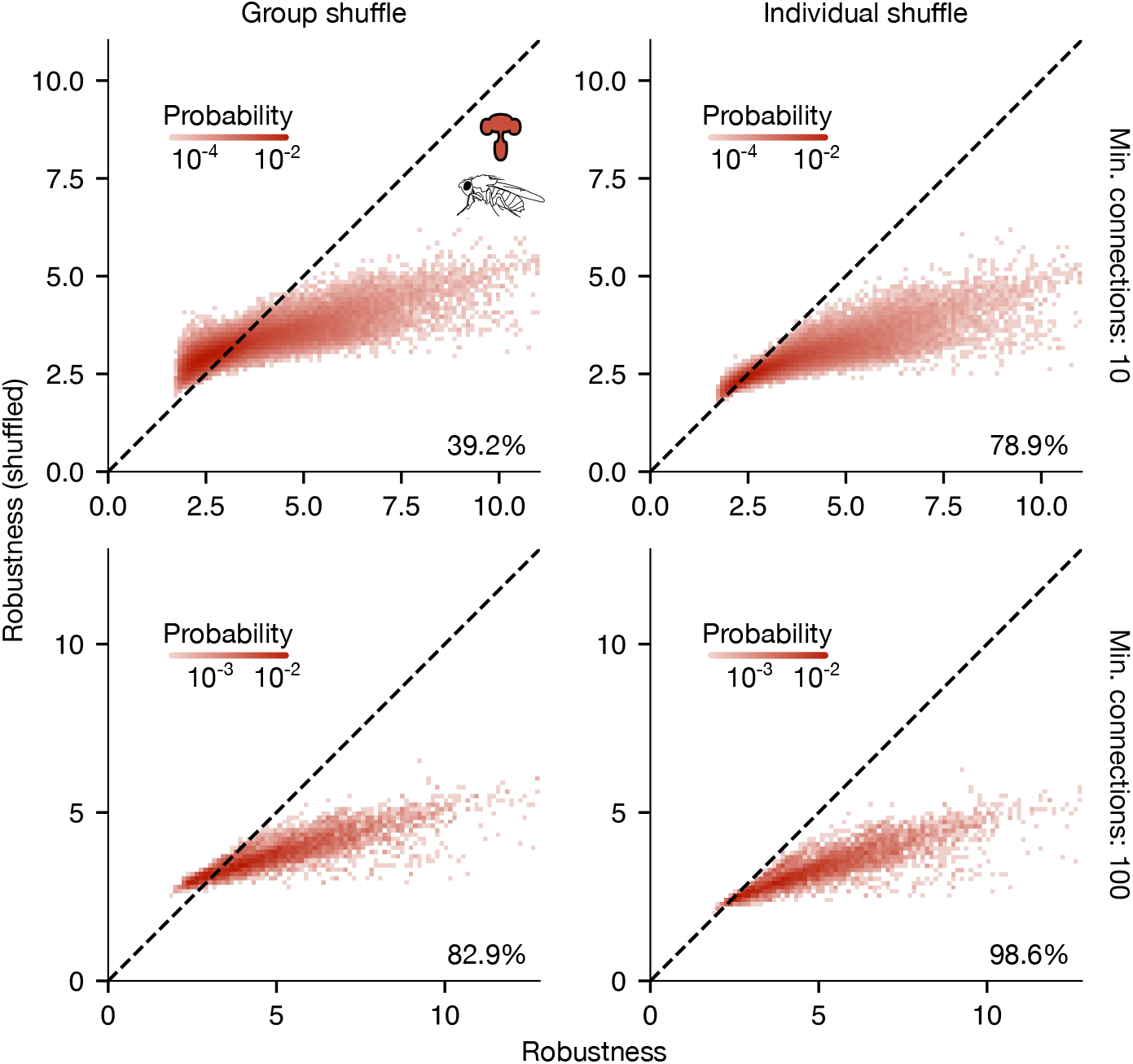
Robustness under different thresholding choices. Robustness of neurons in the fruit fly brain and nerve cord^9^ before and after shuffling synapses among incoming connections. *Left column*: Synapses are placed into connections in groups of three. *Right column*: Synapses placed into connections individually. *Top row*: Comparison for neurons with at least ten incoming connections, as in the main text. *Bottom row*: Comparison for neurons with at least 100 incoming connections.

To test whether the low fraction of high-robustness neurons is explained by neurons with few connections, we restrict our analysis to neurons with at least 100 incoming connections (Fig. S6, bottom row). With this restriction, the low-robustness neurons driving this result are removed, such that the majority of the remaining neurons have higher robustness than their shuffled counterparts.

**Fig. S7.**
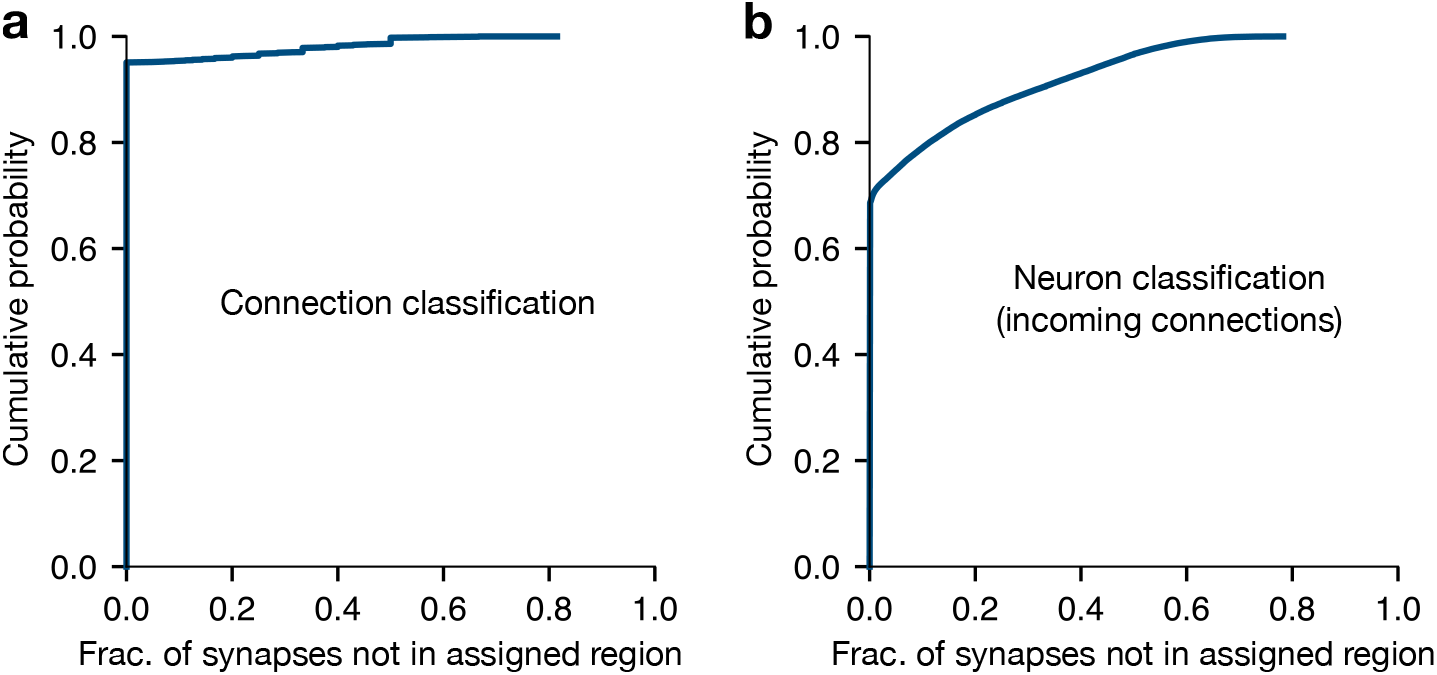
Synapse composition in plurality-classified connections and neurons. **a**, Distribution of the fraction of synapses in each connection that fall outside the connection’s assigned brain region across the adult female fruit fly brain.^7,8^ **b**, Distribution of the fraction of synapses in each neuron’s incoming connections that fall outside the neuron’s assigned brain region. Neurons were classified into brain regions using plurality of their incoming synapses.

### 10. Plurality classification of connections and neurons

Our analysis of synaptic connectivity by brain region in the adult female fruit fly uses a classification of connections and neurons into specific brain regions. Ref. 7, 8 classifies individual synapses into neuropils, and each neuropil belongs to one brain region. We classify connections into brain regions by plurality of their synapses: each connection is assigned the region which contains the largest number of its synapses. Similarly, we assign neurons into brain regions by plurality of their incoming synapses. To quantify how much information is lost in this coarse-graining, we can study the distribution of the fraction of synapses that are in connections or neurons assigned to different regions (Fig. S7). We find that around 95% of connections have all synapses in a single region, and around 70% of neurons have all synapses from their incoming connections in a single region.

